# Cross-species profiling reveals a conserved core of RNA-dependent proteins in yeast

**DOI:** 10.1101/2024.12.06.627129

**Authors:** Nadine Bianca Wäber, Johanna Franziska Seidler, Fabienne Thelen, Palina Kot, Silke Schreiner, Thomas Timm, Vera Bettenworth, Günter Lochnit, Katja Sträßer, Cornelia Kilchert

## Abstract

Delineating the constituents and structural composition of RNA-associated protein complexes is essential to mapping the molecular machinery driving RNA metabolism and its impact on cellular function. Here, we present a comprehensive dataset of RNA-dependent proteins and complexes in the phylogenetically distant yeasts *Saccharomyces cerevisiae* and *Schizosaccharomyces pombe*. Using R-DeeP—a density gradient-based method that uses quantitative mass spectrometry to profile protein sedimentation in the presence and absence of RNA—we introduce an RNA dependence index (RDI) as a descriptive framework for RNA dependence, enabling the robust comparative analysis of RNA dependence across proteins in both species and relative to existing data from their human counterparts. This identifies a conserved core of RNA-dependent proteins shared across both yeasts, alongside distinct, organism-specific adaptations in complex behaviour. The data further support the analysis of co-sedimentation behaviour of protein complexes with known RNA-directed functions. For instance, we find that the five subunits of the *S. cerevisiae* THO complex only co-sediment in the absence of RNA, pointing to an underappreciated structural modularity of the well-characterized pentameric complex. The two datasets, available at https://yeast-r-deep.computational.bio, provide a resource for hypothesis-driven research in RNA biology and establish R-DeeP as a broadly applicable tool for comparative analysis of RNA-protein interactions.

## Introduction

The regulation of gene expression is essential for cellular function, development, and response to environmental cues (1). This regulation operates at all stages of RNA metabolism, including transcription, RNA processing, translation, and degradation, and is largely driven by interactions between RNA and proteins. RNA-binding proteins (RBPs) directly interact with RNA, typically through well-defined RNA-binding domains, to modulate the function, localization, translation, and stability of RNA (2). However, RNA-protein interactions also function outside of RNA metabolism: the activity of protein complexes involved in chromatin regulation, for example, can be modulated through RNA binding (3–5). Over the past decade, advancements in RNA-protein interactomics have substantially expanded the catalogue of RBPs across various organisms, including yeast (6– 17). A preferred method has been RNA interactome capture (RIC), which relies on UV cross-linking of RNA-protein complexes, followed by enrichment of polyadenylated (poly(A)+) RNA and identification of co-purified RBPs by mass spectrometry (MS) (6, 7). RIC has been applied across a wide range of cell types and conditions, with detailed protocols available for various organisms (18–21). Alternative methods rely on either organic phase separation or solid-phase extraction to enrich crosslinked RNA-protein complexes independently of the poly(A) tail (17, 22–25). Combined, these studies identified hundreds of previously unknown RBPs that directly interact with RNA. However, RNA regulation frequently occurs within large, dynamic complexes, where the interaction between proteins and RNA is highly coordinated but not always direct. For instance, certain proteins may play structural or regulatory roles, facilitating the assembly or stability of the entire complex without being in direct contact with RNA themselves (26, 27). Thus, understanding the composition and architecture of RNA-protein complexes, including proteins whose association with RNA is mediated indirectly through other complex members, is critical for uncovering the full spectrum of regulatory mechanisms that govern RNA biology.

Recently, various approaches have been developed to determine RNA-dependent protein interactions; that is, to identify proteins whose complex association or behaviour is contingent on the presence of RNA, whether through direct or indirect contact (28). For example, RBP immunoprecipitation with or without RNase treatment, combined with MS, enables direct comparison of RNA-dependent and -independent RBP interactomes and has been applied to various organisms, including HEK293XT cells at a large scale (27, 29). Other protocols rely on cosedimentation or co-elution to evaluate RNA-dependent macromolecular architecture: Grad-seq combines RNA-sequencing and proteomics of fractionated glycerol gradients to identify comigrating species, enabling the identification of protein complexes associated with specific RNA types such as small RNAs (30–32). DIF-FRAC couples MS to size exclusion chromatography of RNase-treated or untreated reference samples and has been used to characterise human and mouse RNA-protein complexes proteome-wide (33). R-DeeP relies on sucrose density gradient ultracentrifugation to monitor the differential sedimentation behaviour of protein complexes in the presence and absence of RNA. Proteins that shift their sedimentation position upon RNA removal are defined as RNA-dependent; this operational definition, which captures both direct and indirect RNA associations, is adopted throughout this manuscript. R-DeeP has been applied to various human cell types and other organisms including *Plasmodium falciparum* and the cyanobacterium Nostoc sp. PCC 7120, and revealed the RNA association of proteins that had not previously been linked to RNA metabolism (34–38). GradR, a related approach relying on glycerol gradients rather than sucrose gradients, has been employed to identify RBPs in *Salmonella enterica* (39).

Here, we present a proteome-wide atlas of RNA-dependent proteins in the distantly related yeast species *Saccharomyces cerevisiae* (*S. cerevisiae*) and *Schizosaccharomyces pombe* (*S. pombe*), generated using an adapted version of the R-DeeP workflow. To facilitate systematic analysis and cross-experiment comparisons, we introduce the RNA dependence index (RDI), a quantitative score that summarizes the difference in sedimentation behaviour between RNase-treated and control conditions, enabling direct comparison of our yeast datasets with existing human R-DeeP data. Sedimentation profiles of all detected proteins are readily accessible online through a search interface at https://yeast-r-deep. computational.bio/.

Analysis of the resulting dataset revealed several distinct aspects of RNA-dependent complex behaviour. First, comparison across the two yeast species identified a conserved core of RNA-dependent proteins, along with species-specific differences that reflect divergent RNA biology. Second, the sedimentation behaviour of well-characterised ribonucleoprotein complexes (RNPs), including the ribosome and the spliceosome, illustrates how R-DeeP can quantify the degree to which RNA contributes to complex cohesion. Third, for several stable complexes, RNase treatment revealed an unexpected structural modularity, with distinct sub-complexes exhibiting differential RNA sensitivity; detailed analysis of the THO complex suggests that this can reflect RNA-dependent redistribution of individual subunits correlating with different RNA affinities. Finally, only approximately one third of proteins with high RDI scores carried prior annotations linking them to RNA metabolism, underscoring the discovery potential of the approach. Functional enrichment analysis of these unannotated hits revealed a striking and consistent overrepresentation of DNA-binding and chromatin-associated proteins across all three organisms. Using *in vivo* crosslinking, we demonstrate that the high mobility group (HMG)-box chromatin proteins Nhp6a/b (*S. cerevisiae*) and Nhp6 (*S. pombe*) interact directly with RNA, validating R-DeeP as a discovery platform for previously unrecognised RNA-protein associations and establishing HMG-box proteins as a conserved class of RNA-interacting architectural DNA-binding proteins.

## Results and Discussion

### R-DeeP robustly identifies RNA-dependent proteins in budding and fission yeast

We adapted the R-DeeP method (34, 35) to the evolutionarily distant yeasts *S. cerevisiae* and *S. pombe* to compare protein sedimentation during density gradient ultracentrifugation with and without prior RNase treatment to identify RNA-dependent proteins proteome-wide (Figure 1A). After ultracentrifugation, gradients were separated into 25 fractions (Figure S1A and B) and the protein distribution across fractions was analysed by MS. Experiments were carried out with three biological replicates. A total of 2,238 *S. cerevisiae* proteins and 1,991 *S. pombe* proteins were detected across the gradients. For the majority of proteins, sedimentation was unaffected by RNase treatment (Figure 1B and S1C and D), and in these cases, sedimentation patterns were highly consistent across replicates (Figure S1E). Examples for this category are various multiprotein complexes with RNA-unrelated functions (Figure S1F), as reported for the HeLa S3 dataset (34). However, 658 proteins (30.0%) in *S. cerevisiae* and 472 proteins (24.1%) in *S. pombe* did display RNA-dependent shifts (Figure 1B and S1C). For these RNA-dependent proteins, sedimentation patterns were highly reproducible between control replicates but varied substantially across the RNase-treated replicates (Figure S1G). Overall, the proportion of RNA-dependent proteins was lower in both yeasts compared to HeLa S3 cells, where 35.6% of detected proteins shift by at least one fraction after RNase treatment (34) (Figure 1B). Sedimentation profiles as well as raw and normalized fraction data for all *S. cerevisiae* and *S. pombe* proteins are available online at https://yeast-r-deep.computational.bio/.

**Figure 1:**
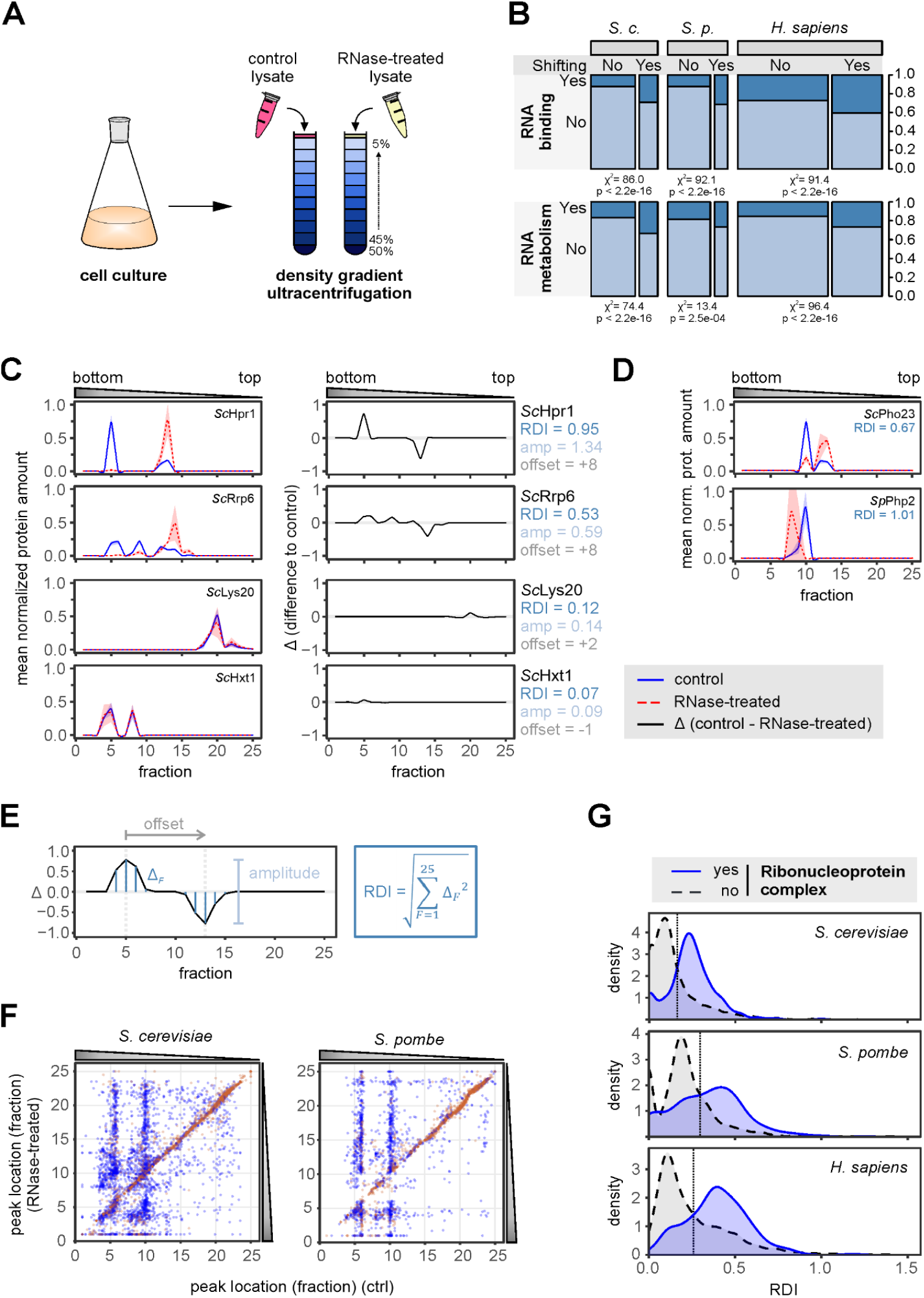
R-DeeP identifies RNA-dependent proteins in yeast. **A** Schematic of the density gradient ultracentrifugation experiment to detect RNA-dependent proteins (R-DeeP). **B** Proportion of proteins among non-shifting and shifting proteins annotated with GO:0003732 “RNA binding” (top) or GO:0016070 “RNA metabolic process” (bottom) for *S. cerevisiae* and *S. pombe* compared to the published human data set (Caudron-Herger et al., 2019). Areas are proportional to protein numbers within each category across organisms. Results of Pearson’s Chi-squared test with Yates’ continuity correction are indicated. Absolute protein counts for each category are provided in Figure S1D. **C** Gradient profiles for two well-characterized RNA-binding proteins in *S. cerevisiae* (Hrp1 and Rrp6; top) and two proteins with no known link to RNA metabolism (Lys20 and Hxt1; bottom). Left: Mean normalized protein amounts across the gradients for control lysates (blue) and RNase-treated lysates (red). The shaded area behind the curves corresponds to +/-one standard deviation. Protein amounts in each fraction were normalized to the total protein amount across the gradient. Right: Difference (Δ, black line) of the normalized protein amounts of the RNase-treated samples relative to the control. The amplitude (amp) and the relative offset (in fractions) of the strongest shift as well as the RNA dependence index (RDI), corresponding to the Euclidean distance between control and RNase-treated curves, were calculated as depicted in E, and are indicated for each protein. **D** Gradient profiles for two RNA-dependent proteins without previous links to RNA metabolism (as in C): *S. cerevisiae* Pho23, component of the Rpd3L histone deacetylase complex (top), and *S. pombe* CCAAT-binding transcription factor complex subunit Php2. **E** Schematic indicating how quantitative indicators for each protein are calculated: amplitude (amp), off-set, and RNA-dependence index (RDI). **F** Positions of protein peaks for the RNase-treated samples relative to the positions in the control gradient for *S. cerevisiae* (left) and *S. pombe* (right). Proteins that peak at a single position in the control gradient are indicated in brown, proteins with multiple peaks in blue. Fractions 6 and 10 correspond to the positions of the large and small ribosomal subunits in the control gradient and contain a large number of shifting proteins. **G** Distribution of RNA dependence indices (RDIs) according to protein annotation as component of a “ribonucleoprotein complex” (GO:1990904) (blue line: yes; grey dashed line: no) in *S. cerevisiae* (top), *S. pombe* (middle), or human (bottom). Human data (HeLa S3 cells) from Caudron-Herger et al., 2019. The vertical hashed line indicates the cut-off used for GO term analysis in Figure 2B.

As expected, shifting proteins in all three organisms were significantly enriched for GO annotations related to RNA metabolic processes, RNA-binding activity, or classical RNA-binding domains compared to non-shifting proteins (Figure 1B, S1C and S1D), validating the approach and providing a baseline against which unexpected hits can be assessed. Representative examples from the *S. cerevisiae* dataset are shown in Figure 1C: Hpr1, a component of the transcription and export (TREX) complex involved in nuclear RNA export (40), and Rrp6, an exoribonuclease associated with the nuclear RNA exosome (41) are among the shifting proteins (Figure 1C, top left panels). As expected for proteins with no known role in RNA biology, the homocitrate synthase Lys20 and the low-affinity glucose transporter Hxt1 did not shift position in the presence of RNase (Figure 1C, bottom left panels). Analogous results were observed in *S. pombe*, where the well-characterised RNP remodelling ATPase Dbp2 and the 40S ribosomal protein S8 shifted in RNase-treated gradients, as confirmed by both sedimentation profiles and Western blot analysis (Figure S1H and I). Beyond this expected behaviour, a substantial proportion of shifting proteins in either yeast lack annotated roles in RNA metabolism or RNA binding (Figure 1B). For example, *S. cerevisiae* Pho23, a component of the Rpd3L histone deacetylase complex (42), and the CCAAT-binding transcription factor complex subunit Php2 in *S. pombe*, both of which regulate chromatin accessibility, shifted in RNase-treated gradients (Figure 1D), warranting further investigation. The complete list of shifting proteins can be found in Supplementary Table S1.

To enable systematic comparison of RNA-dependent protein behaviour both between individual proteins and across species, we sought to define quantitative descriptors of how RNase treatment affects sedimentation patterns. To this end, we normalized the mean amount of each protein in every fraction to the total amount of the respective protein detected across the gradient, then calculated the difference between RNase-treated and control samples (Figure 1C, right panels). The resulting difference curve has a maximum and a minimum, corresponding to the fractions with the greatest loss and gain in protein signal upon RNase treatment, respectively. We define the span between these two extremes as the amplitude of the shift and the number of fractions separating them as its offset (Figure 1E). A positive offset indicates that protein signal shifts from heavier to lighter fractions in the gradient after RNase treatment; that is, the protein sediments more deeply when RNA is intact. Conversely, a negative offset indicates deeper sedimentation in the absence of RNA, which can reflect protein aggregation (note proteins shifting to the bottom-most fraction #1 in RNase-treated gradients in Figure 1F and S1B). In both yeasts, the majority of proteins exhibited a single prominent peak in the control gradients (69.7% in *S. cerevisiae* and 79.8% in *S. pombe*, brown in Figure 1F and S1K). The remaining proteins displayed more complex sedimentation patterns even in the absence of RNase treatment, which was true for the majority of shifting proteins (Figure S1K), suggesting that many RNA interactions are partial or transient in nature. To quantify changes to these complex sedimentation behaviours, which a single amplitude or offset value may not fully reflect, we calculated the Euclidean distance between the control and RNase-treated sedimentation profiles, defined as the square root of the sum of squared fraction-wise differences across all fractions (Figure 1E). This metric captures the overall difference between the curve shapes observed for each protein, with a value of zero indicating identical sedimentation profiles with and without RNase treatment and larger values reflecting greater dissimilarity; in the following, we refer to this score as the RNA dependence index (RDI) (Figure 1E).

Proteins annotated as components of ribonucleoprotein complexes (RNPs) or with known RNA-binding activity not only showed greater shift amplitudes than proteins without the annotation but also significantly higher RDI scores (Figure 1G and S1L and M), a trend observed consistently across yeast and human datasets (RDIs for human proteins were calculated from the published HeLa S3 cell dataset of Caudron-Herger et al., 2019). Importantly, RDI was a stronger predictor of RNA-binding ability and RNP membership than shift amplitude alone in all three organisms, as reflected in the lower p-values (Figure S1L and M), confirming the utility of this metric.

### R-DeeP identifies a conserved core of RNA-dependent proteins in yeast

To compare RNA-dependent protein behaviour across evolutionary distances, we retrieved *S. cerevisiae* orthologues of *S. pombe* proteins from Ensembl Fungi (43, 44). We identified 1,175 orthologous protein pairs where protein signals could be detected in the gradients of both yeasts. While RDIs showed no global linear correlation between the two species, annotated RNP components and RBPs had higher RDIs in both yeasts (Figure 2A, left panel, and Figure S2A). Notably, the direction of the shift was not uniform, although *S. cerevisiae* RBPs had an increased likelihood to display positive shifts (from lower to higher fractions in the gradient) (Figure S2A, right panel). This behaviour was independent of the specific RNA class (rRNA, mRNA, tRNA, or snoRNA) associated with the protein (Figure S2B). To characterize proteins that belong to the conserved core of RNA-dependent proteins more closely, we conducted a gene ontology (GO) term enrichment analysis (45, 46). For this, we defined RDI cutoff values based on the relative behaviour of the populations of RNP components and non-RNP components in both species (Figure 1G). Proteins with conserved, sizable RNA-dependent shifts were frequently associated with RNA-related GO terms (Figure 2B), including “RNA processing”, “protein-RNA complex assembly”, and “cytoplasmic translation” (Figure S2C and D, top panels). In contrast, components of the endomembrane system, signal transduction, or cytoskeleton organization exhibited low RDIs, with offsets centred around zero (signal transduction and cytoskeleton organization) or more likely to be negative than positive (endomembrane system) (Figure S2C and D, bottom panels); the complete list of conserved RNA-dependent proteins can be found in Supplementary Table S2. Consistent with the human dataset, shared components of RNA-associated complexes frequently showed uniform sedimentation behaviour, with complex members shifting to different positions upon RNase treatment. The PeBoW complex, which links ribo-some biogenesis to the cell cycle (47), and is among the conserved shifting complexes, is one such example: Yeast PeBoW components shifted towards higher gradient fractions upon RNase treatment; at the same time, subunits showed an increased tendency to sediment to the pellet fraction, which was also observed for the human Erb1 homologue (Figure 2C), suggesting that the complex has a propensity to aggregate in the absence of RNA. We conclude that the RDI serves as a meaningful quantitative approximation of the degree of RNA dependency and can be used to identify a conserved core of RNA-dependent proteins across species. We also note that for some proteins, such as Pe-BoW components, RNA dependency may not only reflect a shift in sedimentation behaviour but also a tendency towards aggregation upon RNA removal.

**Figure 2:**
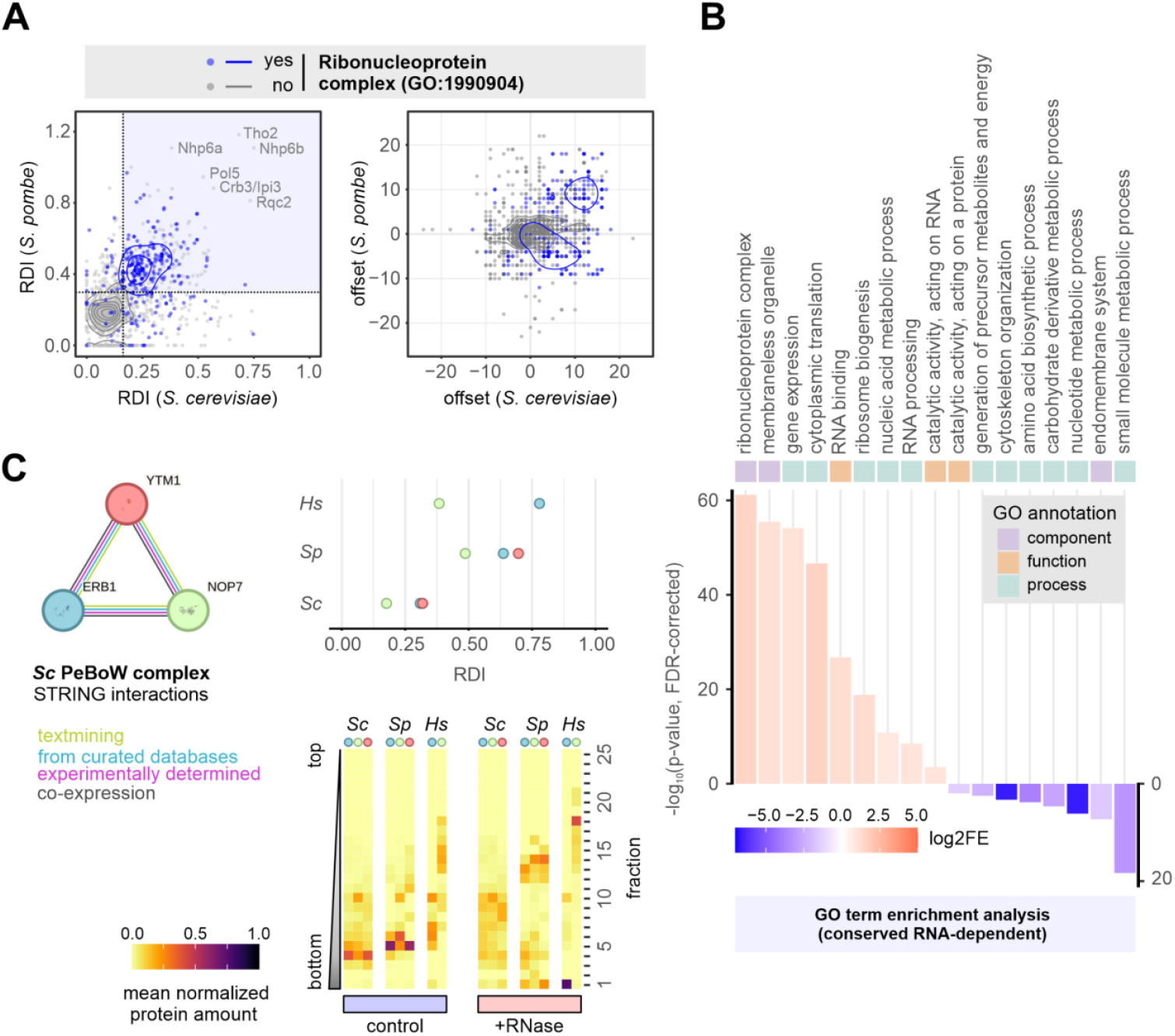
R-DeeP captures a conserved core of RNA-dependent proteins in yeast. **A** Comparison of RNA dependence indices (RDI, left panel) and offset (right) of orthologous protein pairs in *S. cerevisiae* and *S. pombe*. Proteins annotated as “ribonucleoprotein complex” components (GO: 1990904) are shown in blue, all other proteins in grey. Contours of the 2d density estimates for both populations are shown in blue and grey, respectively. The identities of the proteins with the highest RDI in both organisms are indicated. The dotted lines represent the cut-offs used for GO term enrichment analysis in B. **B** GO term enrichment analysis for orthologous protein pairs with high RDIs in both *S. cerevisiae* and *S. pombe* based on the PANTHER classification system and using *S. pombe* identifiers. Colour indicates log2-fold enrichment (log2FE) of proteins with high RDI in both yeasts within each GO term relative to the proportion expected by chance given all members of orthologous protein pairs detected in the experiment (blue: underrepresented, red: overrepresented). P-values were calculated using Fisher’s exact test with false discovery rate (FDR) correction. **C** The PeBoW complex is an RNA-dependent complex in all three organisms. STRING interaction diagram for the PeBoW complex of *S. cerevisiae* (Sc) (left). RDIs for PeBoW complex subunits of *S. cerevisiae* (Sc), *S. pombe* (Sp) and human (Hs) (top right); colours correspond to the Sc orthologues as shown on the left. Sedimentation behaviour in control and RNase-treated gradients (bottom). Human data from Caudron-Herger et al., 2019; the orthologue of Ytm1 was not detected in the human dataset.

### Large RNP complexes are overrepresented in R-DeeP relative to crosslinking techniques

To evaluate strengths and weaknesses of the R-DeeP method relative to classic RNA interac-tome capture (RIC), we first compared RDI values obtained in R-DeeP to the enrichment scores generated by RIC using the *S. pombe* dataset (15). Whereas RIC identifies proteins that directly contact poly(A)+ RNA through UV crosslinking and selective enrichment, R-DeeP detects proteins whose sedimentation behaviour changes upon RNA removal, capturing both direct and indirect RNA associations; these distinct underlying principles suggest that both datasets are complementary rather than redundant, and that a direct comparison may allow to assess what information each method uniquely contributes. Of the proteins detected in the *S. pombe* dataset, 1,848 proteins were identified by both methods (Figure 3A). Consistent with their different designs, the two approaches indeed had distinct predictive strengths: While RDIs more accurately predicted whether a protein belongs to an RNP complex (as indicated by lower p-values), high RIC enrichment scores were a stronger indicator of direct RNA binding (Figure S3A and B). To assess whether these different and potentially complementary strengths also cause the two approaches to preferentially capture different biological processes, we next performed GO term enrichment analysis on proteins that had shown i) high RDI scores in our R-DeeP assays but low enrichment in RIC, i.e., corresponding to the area highlighted in yellow in Figure 3B; or ii) low RDI values in R-DeeP but high enrichment in RIC, corresponding to the area highlighted in blue in Figure 3B. The analysis revealed a clear distinction: GO terms related to processes that involve large, stable RNP assemblies, such as ribo-somes, or large RNA-associated protein complexes, e.g., RNA polymerase (RNAP) complexes and associated factors, were more commonly associated with high RDIs but low RIC scores (Figure 3C and D), consistent with the idea that many of their components associate with RNA indirectly, through their membership of a larger RNA-scaffolded or -associated complex. Conversely, pathways involving transient RNA associations—mainly mRNA processing activities like splicing, 3’-end processing, and RNA decay—exhibited high RIC enrichment but low RDI values (Figure S3C), reflecting their direct contact with poly(A)+ RNA. Smaller stable RNPs, as illustrated above for the trimeric PeBoW complex, frequently received high scores using both approaches (Figure S3A).

**Figure 3:**
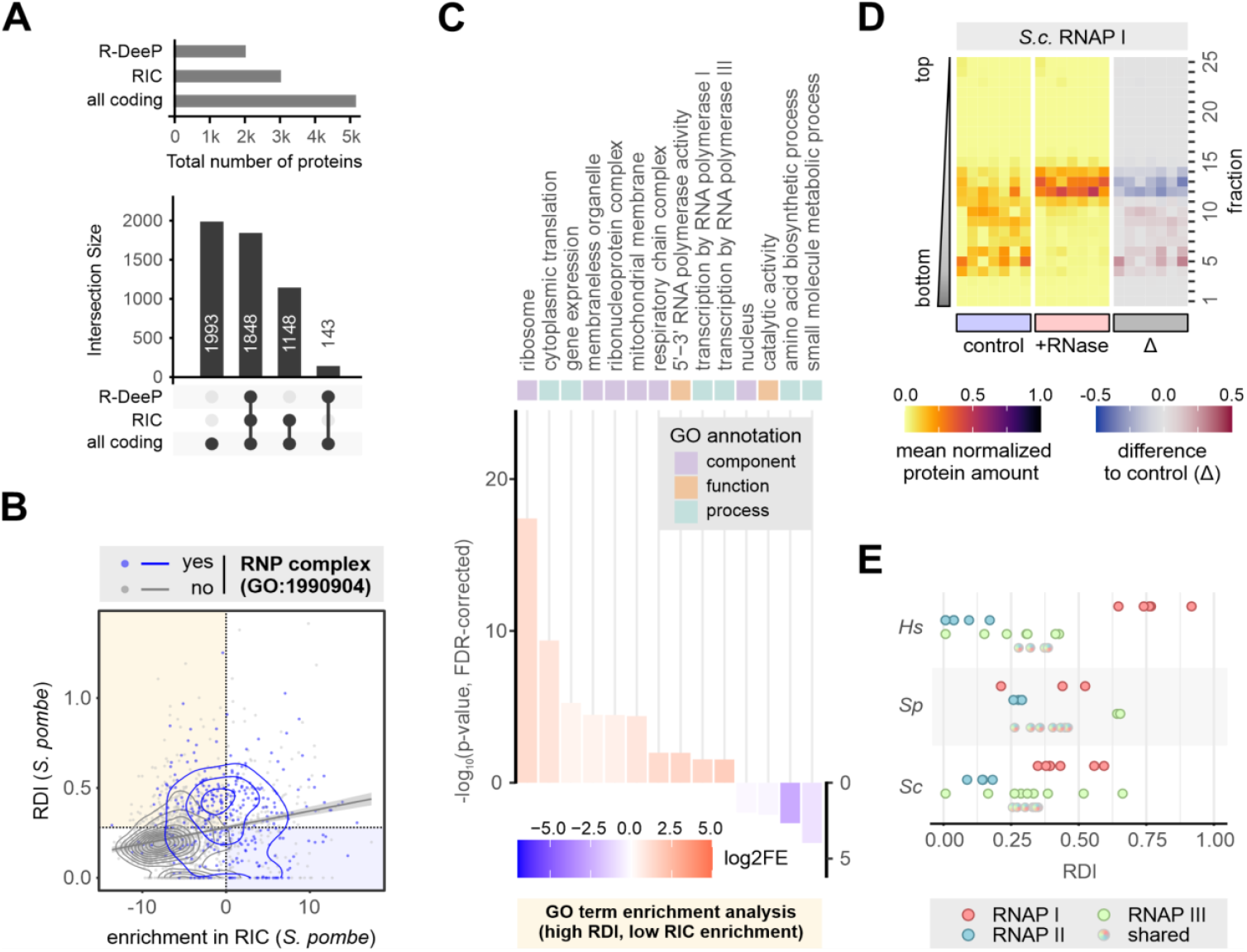
Ribonucleoprotein complexes are overrepresented in R-DeeP. **A** Data availability for *S. pombe* proteins based on the following methods: Density gradient ultracentrifugation (R-DeeP, this study) and poly(A)+ RNA interactome capture (RIC, Kilchert et al. 2020) in comparison to all annotated protein-coding genes (PomBase) **B** Comparison of the RNA dependence index based on R-DeeP (this study) and enrichment after poly(A)+ RNA interactome capture as a measure of direct RNA association (RIC, Kilchert et al. 2020). RIC enrichment is shown as the fold change of mean MS intensities (log_2_) of proteins recovered from oligo(dT) pull-downs of UV-crosslinked samples relative to the whole-cell extract. Proteins annotated as “RNA binding” (GO:0003723) are shown in blue, all other proteins in grey. **C** GO term enrichment analysis for proteins with high RDI and low enrichment in RIC based on the PANTHER classification system. Colour indicates log2-fold enrichment (log2FE) of proteins within each GO term relative to the random distribution of terms across the full dataset (blue: underrepresented, red: overrepresented). P-values were calculated using Fisher’s exact test with false discovery rate (FDR) correction. **D** Sedimentation behaviour of all detected RNA polymerase I components of *S. cerevisiae* in control gradients (left) and RNase-treated gradients (middle). The difference of the mean normalized protein amounts across the RNase-treated gradients to the control is shown on the right, with signals lost after RNase treatment in red and signals gained in blue. **E** RDIs for RNA polymerase (RNAP) subunits of *S. cerevisiae* (Sc), *S. pombe* (Sp) and human (Hs). Colour indicates RNAP type; shared subunits are plotted separately. Human data from Caudron-Herger et al., 2019.

Notably, the subunits of larger RNP complexes tend to cluster at similar RDI values, reflecting a shared degree of RNA dependency within each complex. This clustering is well illustrated by the RNA polymerases: in all three organisms, the subunits specific to each RNAP form distinct clusters, with RNAP II consistently showing the lowest RDI values across species (Figure 3D, E and S3D, E). However, the specific range at which a given complex clusters is not conserved, and the same complex can occupy markedly different RDI ranges in different organisms. In humans, for instance, RNAP I subunits show high RDI values while RNAP III subunits are low-to-medium; in *S. pombe*, the pattern is reversed, with RNAP III subunits showing the highest RDI values and RNAP I only medium values. In *S. cerevisiae*, both RNAP I and RNAP III occupy intermediate ranges. Taken together, these results highlight the complementary nature of R-DeeP and RIC as tools for characterising RNA-protein interactions, and suggest that R-DeeP is particularly effective in capturing proteins whose behaviour is dependent on RNA as a scaffold or structural component of large, stable RNPs, a category that is systematically underrepresented in direct crosslinking-based approaches.

### Structural insights into RNA-scaffolded assemblies

Proteins linked to the ribosome – the largest RNP complex in the cell – constitute the largest subgroup of RNA-dependent proteins, and a very high proportion of proteins involved in translation has high RDIs (Figure 2B and 3C). When considered individually, ribosomal proteins of the large subunit (RPLs) and the small subunit (RPSs) have similar RDIs but vary in the characteristic offset of their shift (Figure 4A and B). Due to their smaller size, subunits of the mitochondrial ribosome (mRP) show distinct sedimentation patterns (Figure 4C, green labels). mRPs were also more resistant to RNase treatment than their cytoplasmic counterparts (Figure 4C, right panel, and S4A and B). In both yeasts, RPLs consistently peaked in fraction #6. RPSs were mainly observed in fraction #10 but were also detected in fraction #6 (Figure 4C and D). This is in contrast to the published HeLa S3 dataset, where protein constituents of both the large and the small ribosomal subunit peak in the same fraction, suggesting the presence of monosomes in the human preparation (Figure S4C) (34). The characteristic sedimentation patterns of RPLs, RPSs, and mRPs in untreated gradients demonstrate that the R-DeeP workflow preserves the integrity of large RNP complexes. Upon RNase treatment, cytosolic RPs dispersed across the gradient, reflecting the disassembly of ribosomal subunits in the absence of stabilising RNA. This illustrates a key feature of the R-DeeP method: its ability to demonstrate the coordinated association of all subunits of an RNA-dependent complex with RNA, thereby generating an inventory of the complex’s protein composition, including components that do not directly contact RNA but are stably incorporated into the same assembly. In contrast, the RIC method captures individual RPs that directly interact with mRNA, for example those forming the mRNA channel of the ribosome (e.g., S3 and S5 in Figure 3D) (15), offering a more detailed view of the suborganization within ribosomal complexes but limited ability to identify the full set of associated components.

**Figure 4:**
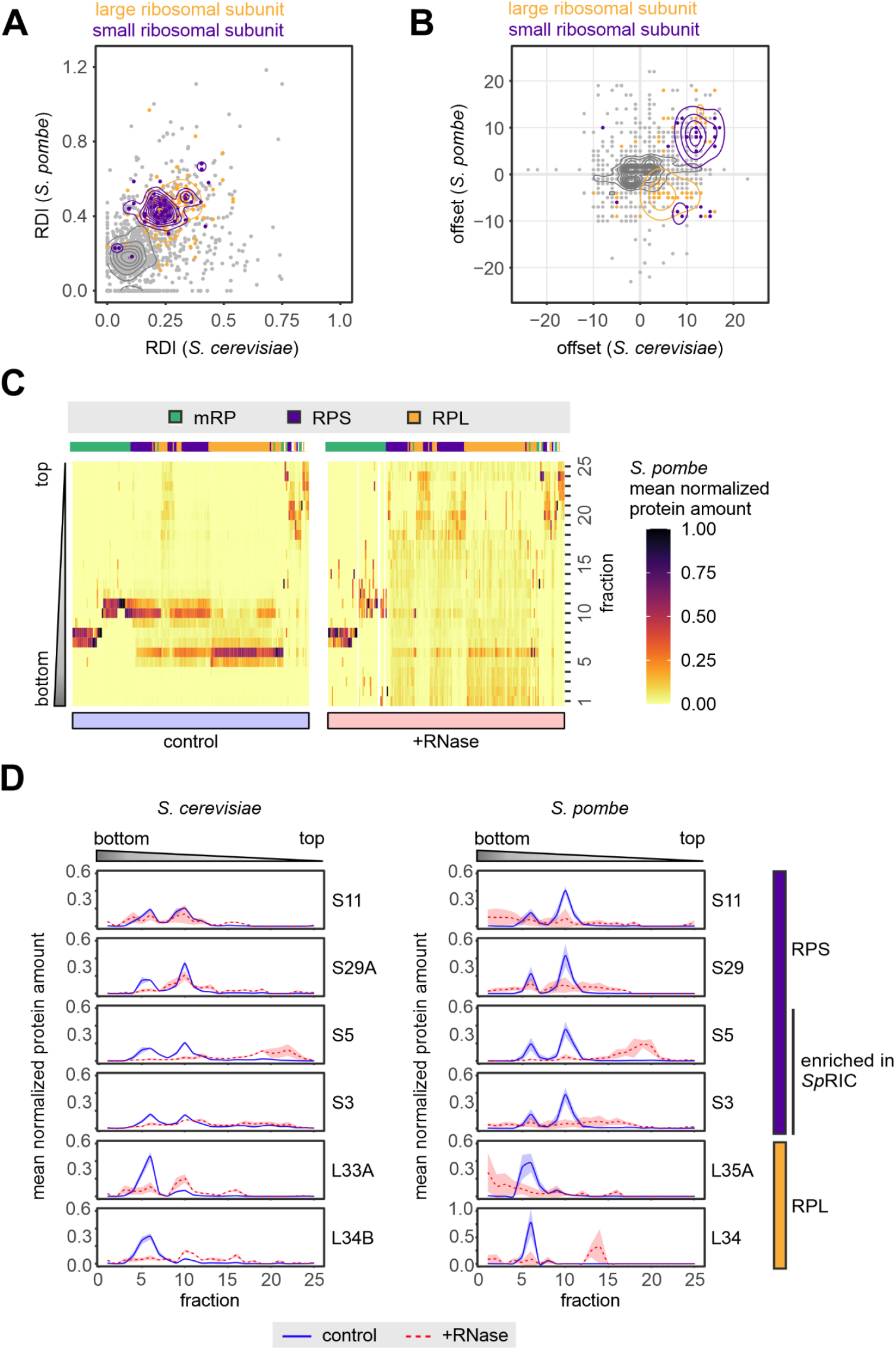
The ribosome as an example of an RNA-scaffolded assembly that disperses after RNase treatment. **A** and **B** Comparison of RDI (A) and shift offset (B) of orthologous protein pairs in *S. cerevisiae* and *S. pombe*. Protein components of the large (GO: 0022625) and small ribosomal subunit (GO:0022627) are shown in yellow and purple, all other proteins in grey. Contours of the 2d density estimates for all populations are shown in yellow, purple and grey, respectively. Annotations were retrieved for the *S. pombe* orthologues. **C** Distribution of *S. pombe* ribosomal proteins in control gradients (left) and RNase-treated gradients (right). Proteins were grouped into 15 clusters identified by hierarchical clustering of all ribosomal proteins (GO:0005840) based on mean normalized protein amounts within all fractions of the control gradients. Both the order of clusters and the order of proteins within clusters were adjusted for clarity of visualization. Proteins of the mitochondrial ribosome (mRP) and the small (RPS) and large (RPL) subunits of the cytoplasmic ribosome tend to cluster together. Annotations are indicated above the plot in green, purple and yellow, respectively. **D** Distribution of different ribosomal proteins across the gradients for *S. cerevisiae* and *S. pombe*. Mean normalized protein amounts in the control and RNase-treated gradients are shown as solid blue and dashed red lines, respectively. The shaded area behind the curves corresponds to +/-one standard deviation. S3 and S5 are adjacent to the mRNA tunnel in the small ribosomal subunit and were highly enriched in the published *S. pombe* RIC (Kilchert et al. 2020).

The spliceosome presents an instructive contrast to the ribosome in this regard. Rather than a stable, stoichiometrically defined RNP, the spliceosome is a highly dynamic assembly that undergoes substantial compositional and structural rearrangements during the splicing reaction and is characterised by a higher protein-to-RNA ratio (48). This raises the question of how such dynamic RNA-dependent assemblies are captured by R-DeeP relative to more stable RNPs like the ribosome. In *S. pombe*, hierarchical clustering tended to group sedimentation profiles of splicing factors associated with specific complexes together; strikingly, members of the Prp19 complex co-migrated as a discrete unit, which sedimented deep into the gradient (fraction #8), an observation not replicated in *S. cerevisiae* or human cells, suggesting that this complex exists as a particularly stable or homogeneous assembly specifically in fission yeast (Figure S4D-F). The Sm core complex, which stably associates with spliceosomal snRNAs, exhibited a more complex sedimentation behaviour (Figure S4D, bottom panel). After RNase treatment, most splicing-related proteins in both yeasts displayed moderate shifts (mean RDI 0.24 and 0.26 in *S. pombe* and *S. cerevisiae*, respectively) with small positive offsets, suggesting that subcomplexes shift as largely intact assemblies and implying that protein-protein contacts are sufficient to maintain cohesion even in the absence of RNA (Figure S4D and E). In contrast, human splicing factors not only exhibited more pronounced shifts (mean RDI 0.39) and higher offsets but also showed a greater tendency to disperse across the gradient upon RNase treatment (Figure S4F), suggesting that RNA plays a more central role in maintaining their integrity. Although the GO term retrieves a considerably larger and more heterogeneous set of proteins in humans than in either yeast, encompassing not only core spliceosome components but also numerous auxiliary factors involved in splice site selection and alternative splicing regulation, the effect extends to core spliceosome components including snRNP proteins (Figure S4F). This may reflect greater spliceosomal complexity and a more extensive dependence on RNA-protein interactions. More broadly, these data illustrate that the pattern of RNase-induced sedimentation changes, i.e., whether a complex shifts as a cohesive unit or disperses, is itself informative about the relative contribution of RNA to the structural integrity of a given RNP assembly.

### Modular sedimentation of RNA-binding complexes in the presence of RNA

The ribosomal and spliceosomal data illustrate how R-DeeP can inform on the degree to which RNA contributes to the cohesion of a complex. However, examining the full dataset, we noticed a puzzling phenomenon: Several well-characterised RNA-binding complexes did not co-migrate as intact assemblies in the presence of RNA. Instead, their subunits distributed across multiple fractions, with co-sedimentation only emerging after RNase treatment (Figure 5A). This was unexpected: if RNA stabilizes these complexes, its removal should cause disassembly, yet in these cases, we observed the opposite. One interpretation is that RNA actively competes with or disrupts the proteinprotein interfaces that hold submodules together, such that intact complexes only form once RNA is removed. To investigate this possibility, we examined the THO complex in detail. The THO complex is a conserved submodule of the transcription and export (TREX) complex that couples RNAPII transcription to nuclear export by packaging nascent RNA and recruiting export factors (40, 49). In *S. cerevisiae*, the pentameric THO core complex (Tho2, Hpr1, Thp2, Mft1 and Tex1) is highly stable, remaining intact during purification under high salt conditions and resisting RNase treatment (50–53). Structural studies further indicate that the complex can dimerize via a specific interface (54). Despite this documented biochemical stability, the *S. cerevisiae* THO complex did not migrate as a single unit in our control gradients. Instead, the complex partitioned into two distinct modules: An RNA-dependent module (consisting of Hpr1, Tho2 and Tex1), which initially migrated in fraction #5 but shifted to fraction #13 after RNase treatment, and the dimerization module (Mft1 and Thp2), identified by structural studies as the primary interaction surface for THO dimerization (54), which sedimented in fraction #13 regardless of RNase treatment (Figure 5B and C). These observations are consistent with the idea that the association between these two submodules is sensitive to the presence of RNA. A potential mechanism involves competition between nucleic acids and negatively charged intrinsically disordered regions (IDRs) for shared binding surfaces within the complex (55). Three THO subunits (Mft1, Hpr1, and Thp2) contain acidic IDRs with isoelectric points ranging from pH 3.8 to 4.0 (56); of these, the IDR of Mft1 is by far the most consequential: with a predicted net charge of -44 at neutral pH, it carries a far greater density of negatively charged residues and thus a higher potential to compete for positively charged RNA-binding surfaces. Notably, this region is absent in the *S. pombe* and human orthologues (Tho7 / THOC7) and is not resolved in the existing crystal structure of the *S. cerevisiae* complex (54). Interestingly, while yeast THO components largely co-sedimented in the RNase-treated gradients, the human complex appeared to lose its integrity upon RNase treatment, with its subunits distributed to different positions in the gradient (Figure 5A) (34).

**Figure 5:**
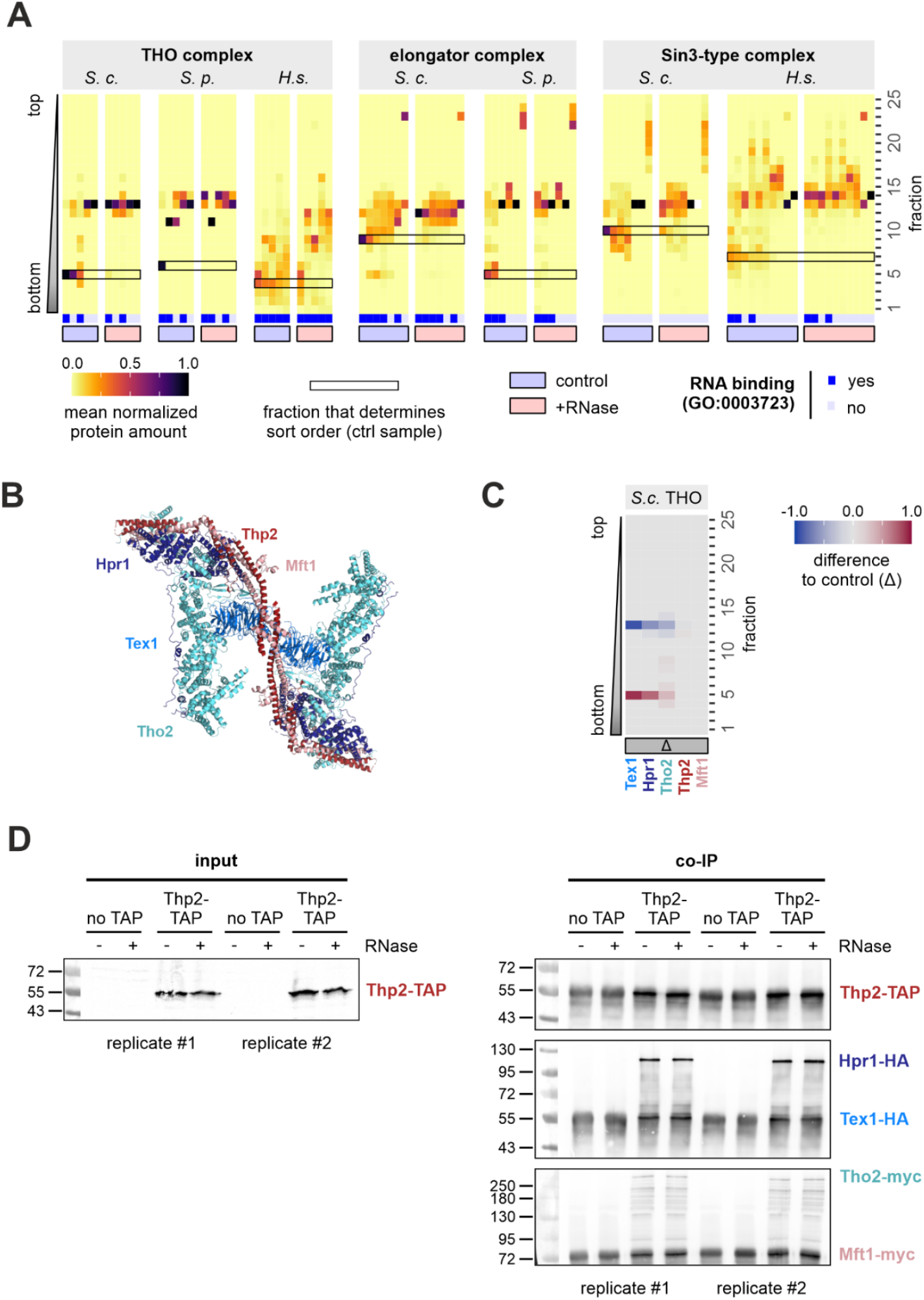
Structural modularity and sedimentation behaviour of multi-subunit RNP complexes. **A** Distribution of THO complex, elongator, or Sin3-type complex subunits in control and RNase-treated gradients. Order of proteins was determined by the signal in the fraction with the partial RNA-dependent signal (indicated). If too few annotated proteins were detected for a given complex in any of the organisms, the data was not included. Annotation of complex components as RNA binding is indicated in blue below the heatmap. Human data from Caudron-Herger et al. 2019. **B** Structure of the *S. cerevisiae* THO dimer (PDB:7AQO) (Schuller et al., 2020) including Hpr1 (dark blue), Tex1 (light blue), Tho2 (turquoise), Thp2 (red) and Mft1 (pink). The structure highlights the importance of Thp2 and Mft1 for the dimerization of the THO complex. The long acidic IDR of Mft1 is not resolved. **C** Redistribution of THO complex subunits of S. cerevisiae after RNase treatment shown as difference of the mean normalized protein amounts across the RNase-treated gradients to the control, with signals lost after RNase treatment in red and signals gained in blue. Colouring of THO subunit labels reflects that in the structure shown in B. Hpr1, Tex1, and Tho2 are part of an RNA-dependent THO module that exhibits a high-amplitude shift. **D** Co-immunoprecipitation of Thp2 from an *S. cerevisiae* strain where all components of the THO complex are epitope-tagged. A strain without tagged Thp2-TAP is included as control. Treatment of the lysates with RNases prior to the purification had no impact on copurification of THO complex subunits. Note that the background signal at the height of Thp2-TAP and Tex1-HA derives from immunoglobulin chains.

To test whether RNA indeed weakens the interaction between the Hpr1-Tho2-Tex1 and Mft1-Thp2 submodules directly, we performed coimmunoprecipitation experiments under the same RNase treatment conditions used in the gradient experiments. However, these batch-based assays did not reveal clear stoichiometric differences in pull-down efficiency that would independently confirm the existence of a stable Hpr1-Tho2-Tex1 subcomplex in the presence of RNA (Figure 5D). This negative result prompted us to consider alternative explanations for the modular sedimentation pattern. Returning to the full dataset, we noted that, within the same complexes, subunits previously annotated as RBPs tended to sediment deeper into the gradient and exhibit more pronounced RNase-induced shifts than their non-RBP counterparts (Figure 5A). This pattern suggests that, in the presence of total RNA, specific subunits may be recruited into interactions with highly abundant transcripts or RNA-rich high-molecular weight assemblies (such as ribosomes), effectively being “dragged” into denser fractions and physically separated from the other complex subunits. Importantly, in pull-down experiments, such spatial separation would be counteracted by continuous mixing of lysate and beads These two mechanisms - RNA-mediated disruption of protein-protein interfaces and RNA-dependent dragging of individual sub-complexes - are not mutually exclusive and distinguishing them for any given complex will require further experimental investigation. Nevertheless, these observations illustrate the potential of R-DeeP to reveal unexpected modularity within well-characterised complexes and to generate specific, testable hypotheses about the role of RNA in regulating their assembly.

### RNA association of chromatin-associated proteins is frequent but warrants *in vivo* validation

In our analysis, only approximately one third of the proteins exhibiting significant RDI scores had previous functional links to RNA metabolism based on existing annotations (Figure 1B). This discrepancy, combined with the method’s particular strength in identifying shifts within medium-to-large complexes, prompted us to investigate whether R-DeeP could identify RNA-dependent complexes that have escaped previ-ous classification. To this end, we performed a functional enrichment analysis using g:profiler (57) to identify protein complexes associated with high-RDI proteins. In addition to the expected enrichment of canonical RNP complexes, there was a striking prevalence of DNA-binding proteins and chromatin-associated complexes. These included replication factor C-like complexes, involved in loading the DNA sliding clamp PCNA during replication and DNA repair (58), as well as core histones (Figure 6A, B and S5A, B). This trend was consistent across both yeast and human datasets: the enrichment of DNA-binding proteins was significant in all three species, and generally more pronounced than that of chromatin components, which reached significance in *S. cerevisiae* and human but not in *S. pombe* (DNA-binding proteins: *S. cerevisiae* p = 3.1e-09, *S. pombe* p = 7.6e-06, human p = 2.8e-32; chromatin: *S. cerevisiae* p = 0.00015, *S. pombe* p = 0.79, human p = 1.3e-09).

**Figure 6:**
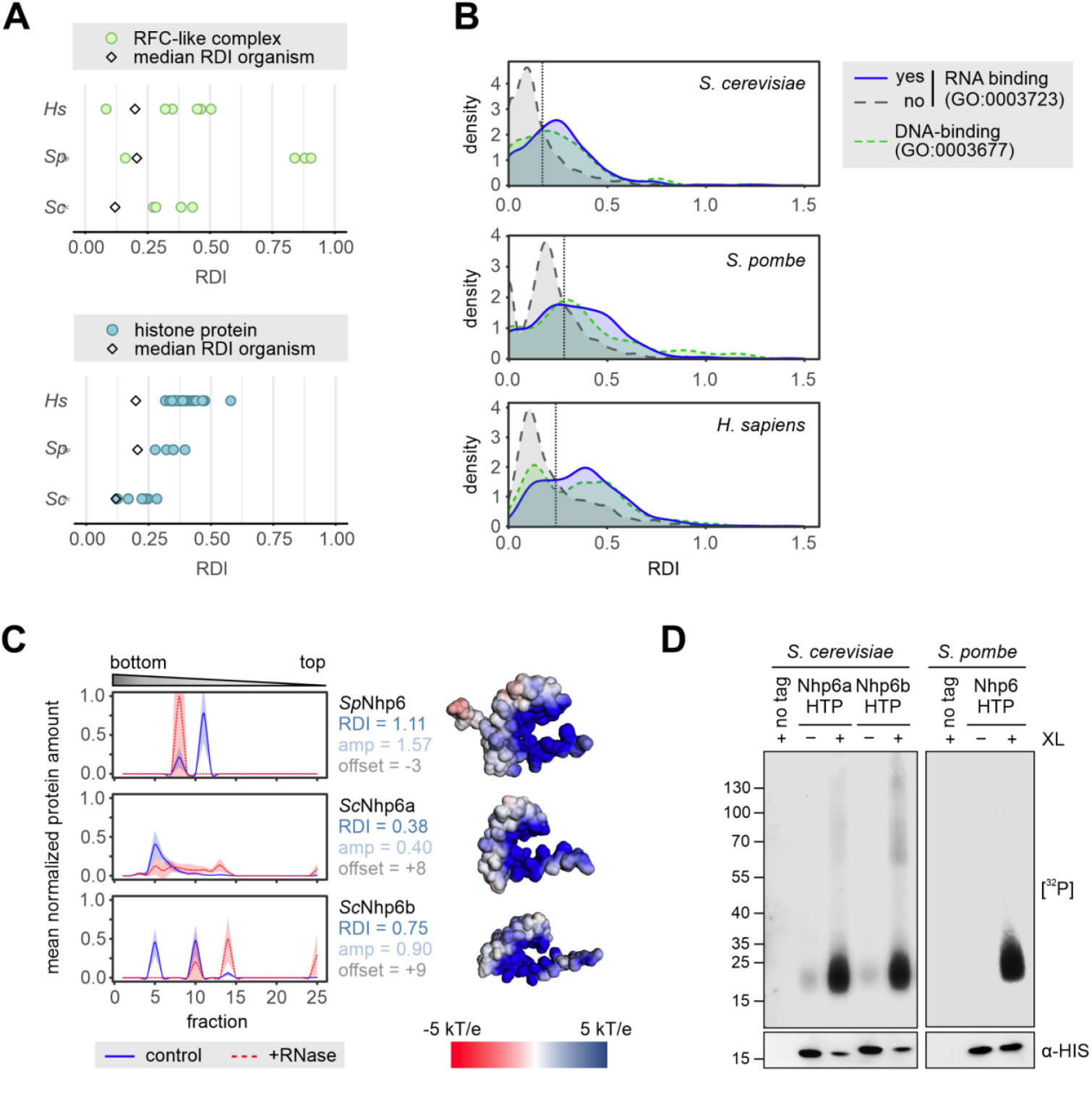
Many DNA-binding proteins display RNA-dependent shifts. **A** RDIs for replication factor C (RFC)-like complex subunits (GO:0031390) (top) and histones (bottom) of *S. cerevisiae* (Sc), *S. pombe* (Sp) and human (Hs). Human data from Caudron-Herger et al., 2019. The diamond marks the median RDI of all proteins detected in the experiment for the given organism. **B** RDI distribution of proteins annotated with GO:0003677 “DNA binding” (green short-dashed line) compared to those annotated or not with GO:0003732 “RNA binding” (blue line vs. grey long-dashed line); human HeLa S3 data from Caudron-Herger et al., 2019. **C** Gradient profiles for HMG-box non-histone chromatin proteins Nhp6 (*S. pombe*), Nhp6a and Nhp6b (*S. cerevisiae*) (left). Mean normalized protein amounts in the control and RNase-treated gradients are shown as solid blue and dashed red lines, respectively. The shaded area behind the curves corresponds to +/-one standard deviation. Electrostatic potential visualisations of Nhp6 Alphafold models (AF-P87057, AF-P11632, and AF-P87057) are shown on the right. **D** Cross-linking and immunoprecipitation analysis of HTP-tagged Nhp6 proteins captured from lysates of UV-crosslinked cells (3J/cm2). After RNase digest, 5′ ends of crosslinked RNAs were radioactively labeled using T4 polynucleotide kinase and [γ-^32^P]ATP, and complexes were separated by gel electrophoresis followed by membrane transfer. The membrane was subsequently probed with anti-HIS-tag antibody to visualise the immunoprecipitated protein. Non-tagged strains were included as controls.

While RNA association has been reported for several individual chromatin-associated complexes and has also emerged more broadly from medium-to large-scale screens, including RNA immunoprecipitation-based approaches and, to a lesser extent, RIC (7, 15, 59), the high frequency of these hits in our datasets raised important questions regarding specificity. A degree of RNA affinity is not unexpected for highly basic proteins such as histones, whose ability to interact with ssRNA has been demonstrated and is driven at least in part through their unstructured tails (60). Substrate promiscuity between RNA and DNA is well documented for various enzyme classes, including ligases and DNA repair enzymes (61, 62). However, the tendency of some nuclear proteins, including histones, to co-migrate with ribosomes in control gradients (Figure S5C) raised the possibility that RNA-dependent sedimentation might reflect adventitious associations that form during lysate preparation rather than native *in vivo* interactions. To determine whether RNA interactions genuinely occur *in vivo*, we employed a crosslinking approach using 4-thiouracil to metabolically label cellular RNA, followed by irradiation at 365 nm to induce zero-distance covalent crosslinks between RNA and interacting proteins. Crosslinked RNA was then detected by radioactive end-labelling with T4 polynucleotide kinase (PNK) following stringent denaturing purification of the protein of interest, an established technique to rigorously validate direct RNA-protein interaction (63).

To distinguish genuine RNA-binding architectural proteins from the more promiscuous, charge-driven associations seen in histones, we focused our validation on the high mobility group (HMG)-box non-histone chromatin proteins Nhp6a/b (*S. cerevisiae*) and Nhp6 (*S. pombe*). Nhp6 proteins ranked among the highest-scoring hits based on their RDI values in both yeasts (Figure 2A and 6C, left panels) and belong to a diverse family of abundant architectural DNA-binding proteins that can bind DNA as monomers or oligomers and participate in a wide range of DNA-dependent processes (64). PNK assays clearly demonstrated that yeast Nhp6 variants are crosslinked to RNA *in vivo* (Figure 6D), providing strong evidence for a direct RNA interaction. Structurally, Nhp6 proteins are small, basic proteins related to human HMGB proteins but containing only a single HMG-box and lacking the pronounced acidic tail found in metazoans, which is involved in autoregulation (Figure 6C, right panels) (65, 66). This experimental confirmation of Nhp6 association with RNA *in vivo* aligns with the recently proposed general capacity of HMG-box domain proteins to interact with RNA (67) and demonstrates that this capacity is conserved beyond the metazoan lineage. Together, these findings demonstrate that RNA-dependent sedimentation of chromatin-associated factors can reflect genuine *in vivo* interactions, while reinforcing that independent experimental verification remains essential to rule out non-specific or artefactual associations.

### Limitations of the method

The analyses presented above demonstrate that R-DeeP can reveal unexpected RNA-dependencies across a broad range of protein complexes, from well-characterised RNPs to chromatin-associated factors with no prior link to RNA biology. As a density gradient-based approach, R-DeeP is relatively straightforward to implement in any laboratory equipped with standard ultra-centrifugation equipment. Compared to UV- or chemical-crosslinking workflows, it requires minimal system-specific optimization, which is primarily limited to the validation of RNase treatment efficiency. Nevertheless, several limitations of the approach deserve explicit mention.

Traditional density gradient analysis is highly sensitive to subtle technical variations, including during gradient set-up, centrifugation, fraction collection, and sample generation, that can shift the absolute fraction in which a protein sediments; the RNase-treated fractions, in particular, are sensitive to variation (compare Figure S1G, H, J). To facilitate quantitative comparisons across experiments despite these variations, we introduced the RDI. By measuring the difference between the control and RNase-treated sedimentation profiles within the same experimental run, the RDI serves as a robust, self-normalizing metric of RNA dependency. This robustness is supported by our observation that subunits of the same stable RNP tend to have similar RDI values. Nonetheless, caution is warranted when comparing RDI values across datasets generated under different experimental conditions.

R-DeeP also shares the inherent limitations common to all MS-based proteomics workflows with limited dynamic range, which can impair detection of low-abundance proteins. This is particularly problematic in gradient fractions that contain the bulk of total cellular protein, particularly the lighter fractions and those containing ribosomal subunits. In these crowded fractions, highly abundant proteins may mask less abundant co-sedimenting proteins, potentially resulting in incomplete sedimentation profiles or false negatives for low-abundance proteins.

In addition, caution must be exercised when interpreting the sedimentation behaviour of proteins with broad nucleic acid affinities. Across all organisms examined, nuclear RNA- and DNA-binding proteins frequently co-sediment with ribosomes, which likely reflects a general electrostatic or structural affinity for nucleic acids rather than native, specific interactions. For proteins and complexes with primarily DNA-directed functions in particular, it is therefore imperative that potential RNA dependencies are validated by orthogonal methods. Crosslinking-based methods such as the PNK assay described above can readily confirm direct RNA-protein interactions; however, validating indirect, RNA-dependent structural associations, where RNA influences complex architecture without being directly bound by the protein in question, remains a significant experimental challenge.

Finally, in its current form, R-DeeP does not involve the biochemical separation of cellular compartments. Consequently, the subcellular localization of the detected proteins or protein complexes cannot be determined from the gradient data alone. In principle, pre-fractionation into cytosolic, nuclear, and membrane fractions could increase spatial resolution and provide valuable biological insights. In practice, however, such steps introduce substantial experimental complexity. Since the number of individual fractions collected for MS analysis scales exponentially (25 fractions per gradient, two treatments (control and RNase), three biological replicates, and multiple cellular compartments or experimental conditions), rapidly reaching several hundred samples per experiment, the associated instrument time and analytical costs must be carefully weighed against the particular suitability of R-DeeP to shed light on the biological question being addressed relative to alternative approaches.

## Material and Methods

### Yeast cultivation

All yeast strains used in this study are listed in Table S3, oligos used for strain generation in Table S4. Standard protocols were used for cell growth and genetic manipulations (68). *S. pombe* and *S. cerevisiae* cells were grown at 30°C in yeast extract with supplements (YES) and YPD (1 % yeast extract, 2 % peptone and 2 % glucose), respectively.

### Protein lysate preparation of *S. pombe*

A 50 mL preculture was used to inoculate 800 mL YES to achieve an optical density (OD) of 0.8 - 1.0 after 15 - 20 h growth at 30°C. Cells were harvested (2000 x g, room temperature (RT), 3 min) and kept on ice. 400 µL lysis buffer (25 mM Tris pH 7.5, 100 mM KCl, 0.02% (v/v) Triton-X, freshly 0.5 mM DTT and 1:10,000 protease inhibitor) were added, and the cells incubated on ice for 10 min. Cells were disrupted with 0.3 g glass beads (Ø 0.1 mm) in a Fastprep-24 5G (MP biomedicals) (four cycles of 1.5 min at 4°C with 15 s pauses between cycles). The cell debris was pelleted (5 min, 5000 rpm, 4°C, followed by max. speed, 15 min, 4°C) and the lysate transferred to a fresh tube. Protein amounts were measured using the bicinchoninic acid assay (BCA Protein Assay Kit, Novagen) on a NanoDrop ND-1000 instrument (Thermo Fisher Scientific). 4 mg of total protein were used for RNase digestion (5 µl RNase Mix (10 µg/mL RNase A, 10 U/mL RNase I, 1000 U/mL RNase T1, 10 U/mL RNase H and 1 U/mL RNase III) per 150 µL lysate) for 1h at 4°C. An equal volume of lysis buffer was added to the control lysate, which was incubated in parallel.

### Protein lysate preparation of *S. cerevisiae*

A 2 L culture was inoculated with an overnight preculture and grown to an OD of 0.8. Cells were harvested at 3600 rpm (Avanti JXN-26 with JLA-8.1) for 3 min at RT and washed with 5 mL ice-cold lysis buffer. Each pellet was resuspended in 500 µL lysis buffer (25 mM Tris pH 7.5, 100 mM KCl, 0.02% (v/v) Triton-X; added freshly: 0.5 mM DTT, 1 µL / 10 mL buffer protease and 1x PhosStop (Roche)). This was followed by cell disruption using a Fastprep-24 5G instrument (MP biomedicals) and 0.3 g glass beads for four cycles of 20 s (6 m/s) at 4°C with 60 s pauses between cycles. Centrifugation of the lysate, determination of protein concentration and RNase treatment was performed as described above.

### Sucrose density gradient preparation

Sucrose density gradients were prepared as described in (35) with the following modifications. Gradients were prepared by layering ten stock solutions, with sucrose concentrations ranging from 50% (w/v) to 5% (w/v) in 100 mM NaCl, 10 mM Tris (pH 7.5), and 1 mM EDTA. The densest solution (3.6 mL) was first placed in a 38.5 mL ultracentrifuge tube (Open-Top Thinwall Polypropylene Tube, 25 × 89mm, Beckman Coulter) and frozen for 15 minutes at -80°C. Subsequently, the remaining sucrose solutions were carefully layered on top, with freezing steps before each addition. Prepared gradients were stored at -20 °C.

### Density gradient ultracentrifugation and fractionation

Six hours before the ultracentrifugation, the frozen sucrose density gradients were placed in the cold room to thaw without disturbing the layers. In the meantime, lysates were prepared as described above. The sample was carefully layered on top of the gradient without touching the surface. Ultracentrifugation was carried out in an Optima XPN-100 Ultracentrifuge (Beckman Coulter) equipped with a SW32 Ti swinging bucket rotor (Beckman Coulter) at 30,000 rpm (acceleration / deceleration set to 5) for 18 h at 4°C. The tubes were carefully removed from the rotor and fractionated bottom-to-top using an ÄKTA Start (Cytiva) with fractionator and set dead volume. At the start, a tube with buffer was carefully placed on the lower part of a Brandel fractionation holder and connected to the sample pump. Now, 10 ml buffer was used to fill the tubing up to the fractionator at a rate of 1 mL/min. Then the buffer tube was carefully exchanged against the first gradient. Fractionation was performed at 1 mL/min with a fraction size of 1.5 mL (25 fractions in total). After the run, the tubing was rinsed with at least 10 mL buffer before fractionating the next gradient. Factions were stored at -80°C or precipitated directly for mass spectrometry or Western blot analysis.

### Protein precipitation

1 mL fraction volume was incubated with 250 µL E-TCA solution (10% trichloroacetic acid, 80% acetone, 0.0015% deoxycholate). The suspension was inverted and stored at - 20°C for at least 1 h or overnight before centrifugation (max. speed, 5 min, 4°C). The protein pellet was washed with 500 µL ice-cold 100% acetone (max. speed, 5 min, 4°C) and air-dried for 5 min. The pellet was stored at -80°C or directly used for mass spectrometry. For Western blot analysis, 60 µl TUS-buffer (50 mL Tris-HCl pH 7.4, 100 mM NaCl, 8 M urea) and 20 µL 4x SDS-loading buffer (100 mM Tris-HCl pH 6.8, 8% (w/v) SDS, 8% (v/v) β-mer-capto-ethanol, 0.04% (w/v) bromophenol blue, 30% glycerol) were added and heated at 95°C for 15 min.

### Sample preparation for TMT-labelling

All chemicals were at least “HPLC grade” or “mass spectrometry grade”. 50 µl 50 mM [NH_4_]_2_CO_3_ buffer were added to the protein pellets. The tubes were vortexed, then placed into an ultrasound bath for 10 min. To pellets that were not fully dissolved, 1 µL ProteaseMax™ Surfacant (Promega) was added, then 50 µL 50 mM [NH_4_]_2_CO_3_ / 5 mM DTT, each followed by two cycles in an ultrasonic bath. For each fraction, the final volume was noted. The following instructions apply to a sample volume of 50 µl; for larger samples, volumes were adjusted accordingly. First, 5 µL 5 mM iodoacetamide (IAA) were added and the samples incubated in the dark for 30 min at RT. To stop the reaction, 3 µl 85 mM cysteine were added and incubated at RT for 15 min. Proteins were digested overnight at 37°C with 2 µg trypsin (Trypsin Gold, Mass spectrometry Grade, Promega). Digestion was stopped by adding 70 µL 1% trifluoroacetic acid (TFA) per 50 µL initial sample volume. After a brief incubation at RT, the sample volume was increased to 500 µL with 0.1% TFA before purification over C18 columns (C18-EC, chromabond, Macherey-Nagel). Columns were washed with 500 µL 0.1% TFA. After sample addition, the columns were washed three times with 500 µL 0.1% TFA, followed by elution with 3 × 200 µL 0.1% TFA / 60% acetonitrile (ACN). Elutions were then pooled. For TMT labelling, the samples were placed in the speed-vac (Concentrator plus, Eppendorf) for 1 h, followed by lyophilisation overnight (ALPHA 1-4 LDC-1M, Christ).

### TMT-labelling

For TMT labelling, 0.8 mg of the label was dissolved in 170 µL 100% ethanol, sufficient for six samples. TMTsixplex™ (Thermo Fisher Scientific) was used for *S. pombe* and TMTpro-18-plex (Thermo Fisher Scientific) for *S. cerevisiae*. The isobaric compounds present in the Reagent Set were designated to the samples as follows: reporter ions at m/z = 126 – Sp-wt-ctrl1, 127 – Sp-wt-ctrl2, 128 – Sp-wt-ctrl3, 129 Sp-wt-rnase1, 130 – Sp-wt-rnase2, 131 – Sp-wt-rnase3 [TMTsixplex™]; 126 – Sc-wt-ctrl1; 127N – Sc-wt-rnase1, 127C – Sc-wt-ctrl2, 128N – Sc-wt-rnase2, 128C – Sc-wt-ctrl3, 129C Sc-wt-rnase3 [TMTpro-18-plex]. The distributing markers TMTpro-129N to 135N were used for samples not related to this study. The lyophilized samples were resuspended in 50 µL 50 mM HEPES, pH 8. 20 µL TMT label was added and incubated for 1 h at RT. The reaction was stopped by addition of 50 µL of 250 mM cysteine and incubated at RT for 15 min. Samples were combined by fraction number and dried overnight in the lyophile (ALPHA 1-4 LDC-1M, Christ). Samples were dissolved in 500 µL 0.1% TFA and purified on C18 columns as before (C18-EC, chromabond, Macherey-Nagel). For this, the columns were primed with 500 µL 0.1% TFA / 60% ACN, then washed three times with 500 µL 0.1% TFA. After sample application, the columns were washed twice with 500 µl 0.1% TFA / 5% methanol before elution with 3 × 200 µL 0.1% TFA / 60% ACN and pooling of the elution fractions. Samples were then placed in the speed-vac (Concentrator plus, Eppendorf) for 1 h and freeze-dried in the lyophile (ALPHA 1-4 LDC-1M, Christ) overnight. Finally, pellets were dissolved in 25 µL 0.1% TFA. To samples that were not fully dissolved, an additional 25 µL 0.1% TFA were added. If that was not sufficient to fully dissolve the pellets, 25 µL 0.1% TFA / 10% ACN was added until all samples were fully dissolved. The final volume of each sample was noted. The protein concentration was determined on a NanoDrop™ 2000 (Thermo Scientific) and samples diluted to 1 µg / µL with 0.1% TFA in a total volume of 15 µL. Samples were stored at -20°C until further use.

### Liquid Chromatography and Tandem Mass Spectrometry (LC-MS/MS/MS)

For MS analysis, 1 µg of each sample was loaded onto a 50 cm µPAC™ C18 column (Pharma Fluidics, Gent, Belgium) in 0.1% formic acid (Fluka Chemie) at 35°C. Peptides were eluted with a linear gradient of ACN from 3% to 44% over 240 min followed by a wash with 72% ACN at a constant flow rate of 300 nL / min (Thermo Fisher Scientific™ UltiMate™ 3000 RSLC nano) and sprayed into an Orbitrap Eclipse Tribrid mass spectrometer (Thermo Fisher Scientific) using an Advion TriVersa NanoMate (Advion BioSciences). The MS was operated in the positive-ionization mode with a spray voltage of the NanoMate system set to 1.7 kV and source temperature at 300°C. Using the data-dependent acquisition mode, the instrument performed full MS scans every 2.5 seconds over a mass range of m/z 400–1600, with the resolution of the Orbitrap set to 120000. The RF lens was set to 30%, auto gain control (AGC) was set to standard with a maximum injection time of 50 ms. In each cycle, the most intense ions (charge state 2-6) above a threshold ion count of 5.000 were selected for CID fragmentation (isolation window of 0.7 m/z) at a normalized collision energy of 35% and an activation time of 10 ms. Fragment ion spectra were acquired in the linear IT with a scan rate set to rapid and mass range to normal and a maximum injection time of 100 ms. After fragmentation, the selected precursor ions were excluded for 20 s from further fragmentation. From each MS/MS cycle, up to ten fragment ions were selected for higher-energy collision-induced dissociation (HCD) fragmentation (isolation window of 3 m/z) at normalized collision energy of 75%. MS3-fragment ion spectra were acquired in the Orbitrap with a resolution of 50000. The mass range was set to 100 – 500 m/z, maximum injection time to 100 ms and AGC to 300.

### Protein identification and quantitation

Data were acquired with Xcalibur 4.3.73.11. (Thermo Fisher Scientific) and the resulting datasets analysed with Proteome Discoverer 2.4.0.305 (Thermo Fisher Scientific). Sequest HT (Proteome Discoverer version 2.4.0.305; Thermo Fisher Scientific) was used to search against the *S. pombe* or *S. cerevisiae* database. A precursor ion mass tolerance of 10 ppm was used, and one missed cleavage was allowed. Carbamidomethylation on cysteines and labelling with TMTpro-18plex at peptide N-termini and lysine side chains were defined as a static modification with optional oxidation of methionine. The fragment ion mass tolerance was set to 0.6 Da for the linear ion trap MS2 detection. The false discovery rate (FDR) for peptide identification was limited to 0.01 by using a decoy database. Protein identifications were accepted if they could be established with at least one unique identified peptide of a length between 6 and 144 amino acids. TMT reporter ion values were quantified from MS3 scans with an integration tolerance of 20 ppm. As samples were pooled by fraction number prior to MS analysis (see TMT labelling section), the TMT reporter ion channel served to assign each quantified peptide to its fraction of origin and experimental condition. Reporter ion intensities were summed per protein across all peptides assigned to each channel. No normalisation across fractions was applied.

### Data analysis and visualization

GO term annotations, Pfam ID annotations, and yeast orthologues were retrieved from Ensembl using biomaRt (43, 44, 69). Additional *S. pombe* sequences and functional annotations were retrieved from PomBase (70). GO term enrichment analysis was carried out based on the PANTHER classification system at https://geneontology.org/ (45, 46). Plots were generated in R Studio with ggplot2 using viridis colour scales and the geom_xspline() function in the ggalt package for spine plots (71–73). Scripts used for hierarchical clustering of ribosomes / spliceosome were based on a published workflow (https://github.com/Tsvanemden/Martin_Caballero_et_al_2021) (74). R code used to generate the figures is available on Github (https://github.com/Kilchertlab/R-DeeP). Electrostatic potential visualisation for Nhp6 proteins was carried out with the APBS-PDB2PQR software suite (v3.4.1 / v3.6.1) based on Alphafold models (75).

### Co-immunoprecipitation

400 mL of *S. cerevisiae* culture were harvested at OD 0.8, the pellet washed with 5 mL PBS and resuspended in 1 mL of IP buffer (25 mM Tris, pH 7.5, 100 mM NaCl, 1 mM EDTA, 0.02% Triton X-100) containing freshly added 0.5 mM DTT, 1x protease inhibitor cocktail (Merck, #P8215-1ML) and PhosSTOP phosphatase inhibitor cocktail (Merck, #4906845001). Cells were lysed in a FastPrep®-24 5G bead beating grinder and lysis system (three 20 sec cycles (6 m/s) with 1 min breaks on ice) using glass beads (500 µL of beads per 750 µL of cell suspension). The lysate, separated from the beads, was centrifuged at 13000 rpm 4°C for 10 min. Supernatants were treated with 10 µL RNase mix (10 µg/ml RNase A, 10 U/ml RNase I, 1000 U/ml RNase T1, 10 U/ml RNase H and 1 U/ml RNase III) or IP buffer for 60 min at 4°C on a turning wheel. 30 µL of each sample were taken as input control. The remaining supernatant was incubated with 30 µL M-280-Tosylactivated Dynabeads (Invitrogen, #14204) coupled to rabbit IgG (ChromPure, #011-000-003) for 18 hr at 4°C on a turning wheel. Beads were washed ten times with ice-cold IP buffer and bound complexes released by TEV cleavage (0.2 mg/mL TEV protease for 2 hr at 16°C on a turning wheel). Samples were analyzed by SDS-PAGE and Western blot (anti-TAP tag polyclonal antibody (Invitrogen, #CAB1001), anti-myc tag polyclonal antibody (Sigma-Aldrich, #SAB4301136), anti-HA tag antibody (Sigma-Aldrich, #H6908).

### Crosslinking and immunoprecipitation

From a YES pre-culture, 1 L cultures were inoculated for overnight growth (*S. cerevisiae*: SDC-lowURA (10 mg / l uracil); *S. pombe*: EMMG-lowURA (10 mg / l uracil)). At an OD600 of 0.5-0.7, 10 ml 4-thiouracil (4tU) solution (10 mg / ml in DMSO; Sigma 440736-1g) were added, and cultures grown for a further 3 h (*S. cerevisiae*) or 4.5 h (*S. pombe*) to allow 4tU incorporation. Cells were harvested by filtration. Non-crosslinked controls were scraped directly into liquid N2 and stored at -80°C until further use. Otherwise, cells were resuspended in 40 ml ice-cold PBS, and UV-irradiated (365 nm; 3 J / cm2) in a 15 cm petri dish on ice in a Stratalinker device. The cell suspension was collected, pooled with 10 ml PBS used to wash remaining cells off the petri dish, and cells frozen in liquid N_2_ after pelleting by centrifugation. Cells were lysed by vortexing after addition of 1 volume TN150 buffer (50 mM Tris-HCl pH 7.8, 150 mM NaCl, 0.1% NP-40, 1 mM DTT (added freshly)) and 2-3 volumes of acid-washed glass beads (G8772, Sigma) for 30min with 30 s on/off intervals on ice. After addition of 3 volumes TN150 buffer / gram cell pellet and vortexing, cellular debris was removed by centrifugation (20 min, 4000 rpm, 4°C). The lysate was cleared by further centrifugation in 2 ml microcentrifuge tubes (20 min, 20,000 x g, 4°C). 2.5% of the sample were removed as input control. Lysates were nutated with 0.2 ml rabbit IgG Sepharose slurry (A2909-5ML, Sigma; pre-washed with TN150 buffer) in 15 ml tubes for 2 h at 4ºC. Beads were washed twice with 5 ml TN1000 buffer (50 mM Tris-HCl pH 7.8, 1 M NaCl, 0.1% NP-40, 1 mM DTT (add freshly)) and twice with 5 ml TN150 buffer (3 min, 1000 rpm, 4ºC), then transferred to a 1.5 ml microcentrifuge tube in a small volume of TN150 buffer. Beads were then resuspended in 240 µl of TN150 buffer (without protease inhibitors and DTT), and 4-8 µl of HaloTEV protease (G6601, Promega) added. TEV cleavage was allowed to proceed for 2h @ 16°C, and the supernatant collected. Volumes were adjusted to 550µl with TN150 buffer, 1µl RNase I (Jena Bioscience, 70u/µl) added and incubated for 5 min at 37°C. 0.4 g of guanidium hydrochloride were then added to the TEV eluates (∼ 0.5 ml of powder; final concentration 6M) and dissolved by vortexing. NaCl was then added to a final concentration of 300 mM (27 µl 5M NaCl stock solution) and imidazole to a final concentration of 10 mM (3 µl 2.5M Imidazole, pH 8.0). TN150 buffer was added to a final volume and the solution nutated with 50 µl of Nickel-NTA beads equilibrated with wash buffer I (50 mM Tris-HCl pH 7.8, 300 mM NaCl 10 mM imidazole, 6 M guanidine hydrochloride, 0.1% NP-40, 1 mM DTT (added freshly)) overnight at 4ºC. The beads were washed twice with 500 µl wash buffer I and three times with 500 µl 1x PNK buffer (50 mM Tris-HCl pH 7.8, 10 mM MgCl2, 0.5% NP-40, 1 mM DTT (added freshly)). Polynucleotide kinase treatment was carried out on-bead by adding 10 µl 1x PNK mix (per sample: 1 µl 10x PNK buffer (NEB), 2.5 µl [γ-^32^P]ATP (SCP-801; Hartmann analytic), 0.5 µl T4 PNK enzyme (M0201S; NEB), 0.25 µl RNaseOUT (Thermo Fisher), 5.75 µl water) and incubating 20 min at 37°C. Beads were washed twice with cold TBS-T, twice with cold 1x PNK buffer, and bound proteins eluted by boiling in 30 µl 1x LDS loading buffer with reducing agent (Invitrogen) for 10 min at 70°C. Proteins were separated on NuPAGE 4-12% Bis-Tris mini-gels (Invitrogen) and transferred to nitrocellulose. Radioactive signal was detected by exposing the membrane to a storage phosphor screen and detected with a Typhoon FLA 9500 Imager (GE Healthcare), followed by antibody-based detection of His-tagged proteins (6x-His Tag Monoclonal Antibody HIS.H8, MA1-21315, Thermo Fisher).

## Data Availability

The mass spectrometry proteomics data have been deposited to the ProteomeXchange Consortium via the PRIDE partner repository (76) with the dataset identifier PXD058607. Gradient profiles can be accessed at https://yeast-rdeep.computational.bio/, where the complete processed datasets are also available for download. R scripts used for data analysis as well as the complete datasets are available on github (https://github.com/Kilchertlab/R-DeeP).

## Acknowledgements and Funding

We thank Maïwen Caudron-Herger and Sven Diederichs for kindly providing their processed human R-DeeP data. The Nab2 antibody was a gift of Maurice Swanson. This work was funded by the Deutsche Forschungsgemeinschaft (DFG) via the Emmy Noether Programme (KI1657/2-1 to CK) and the graduate training group GRK2355 to CK and KS, and by the EU via the ERC Consolidator Grant “mRNP-PackArt” to KS. It was also supported by the de.NBI Cloud within the German Network for Bioinformatics Infrastructure (de.NBI) and ELIXIR-DE (Forschungszentrum Jülich and W-de.NBI-001, W-de.NBI-004, W-de.NBI-008, W-de.NBI-010, W-de.NBI-013, W-de.NBI-014, W-de.NBI-016, W-de.NBI-022) and received technical assistance from the Bioinformatics Core Facility at the professorship of Systems Biology at JLU Giessen. ChatGPT (OpenAI) and Claude Sonnet 4.6 were used for language editing.

## Supplementary Data

**Supplementary Table 3.** Yeast strains used in this study.

| Strain | Organism | Description | Genotype | Reference |
| --- | --- | --- | --- | --- |
| YPCK426 | *S. pombe* | wild type | h+ *leu1-32 ura4-D18 ade6-M216* | Aydin et al., 2024 |
| YPCK100 | *S. pombe* | Dbp2-HTP | h+ *leu1-32 ura4-D18 ade6-M216 dbp2::dbp2-6xhis-TEV-protein A:kanMX* | Aydin et al., 2024 |
| YPCK906 | *S. pombe* | Nhp6-HTP | h+ *leu1-32 ura4-D18 ade6-M216 nhp6::nhp6-6xhis-TEV-proteinA:kanMX* | This study |
| Y1 (RS453) | *S. cerevisiae* | wild type | MAT a *ade2-1 his3-11,15 ura3-52 leu2-3,112 trp1-1 can1-100 GAL+* |  |
| KS3640 | *S. cerevisiae* | THO five tag | MAT a *ura3-1 trp1-1 his3-11,15 leu2-3,112 ade2-1 can1-100 GAL+ Tho2-9myc::KanMX4 Tex1-6HA::Leu Mft1-3myc::Trp Hpr1-6HA::His Thp2-TAP::Ura* | Stefanyshena et al. 2026 |
| YPCK907 | *S. cerevisiae* | Nhp6a-HTP | *met15Δ0 ura3-0 leu2-0 his3-1 NHP6A-6xHIS-TEV-ProtA::kanMX* | This study |
| YPCK908 | *S. cerevisiae* | Nhp6b-HTP | *met15Δ0 ura3-0 leu2-0 his3-1 NHP6B-6xHIS-TEV-ProtA::kanMX* | This study |

**Supplementary Table 4.**
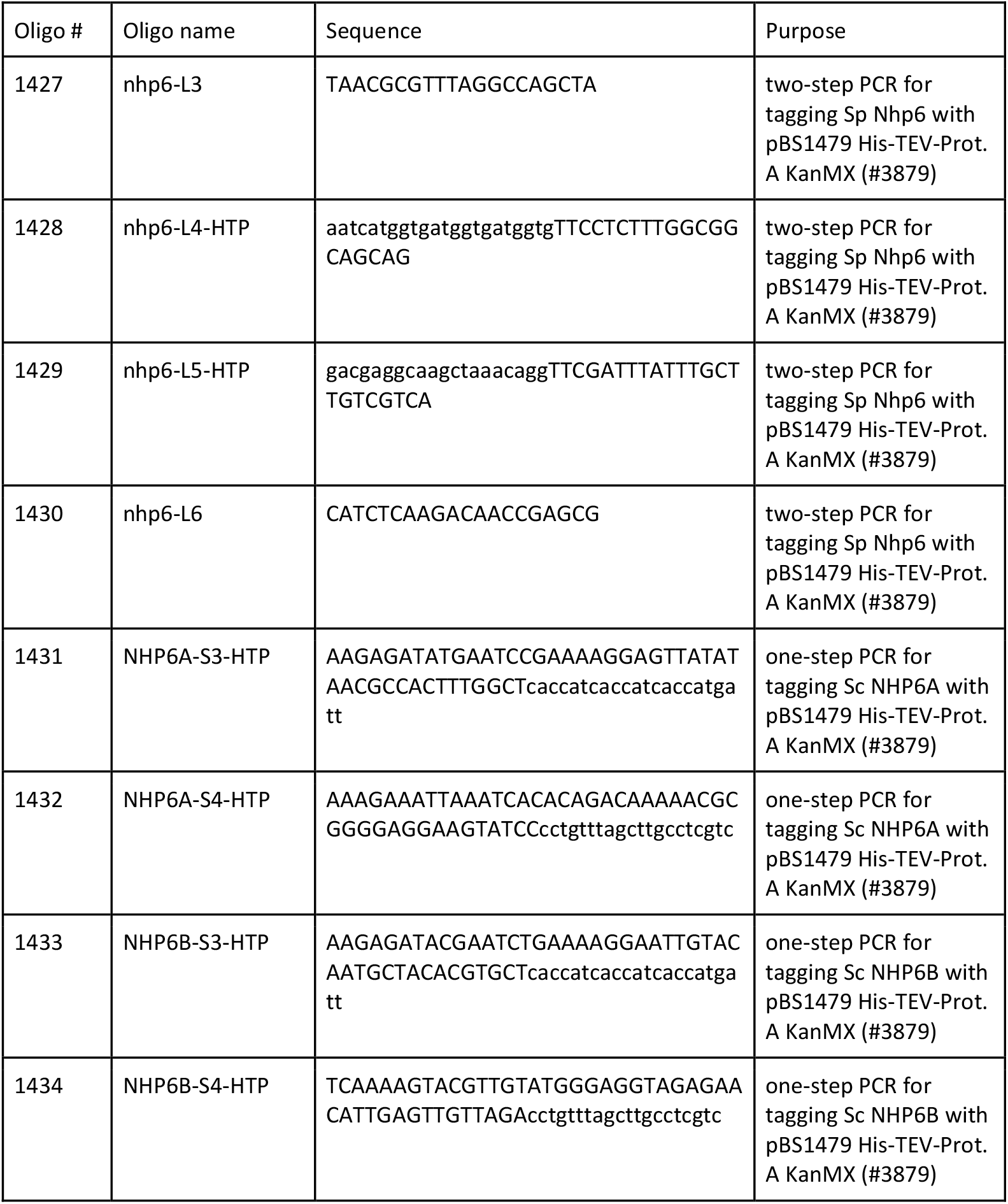
Oligos used in this study.

**Supplementary Figure 1:**
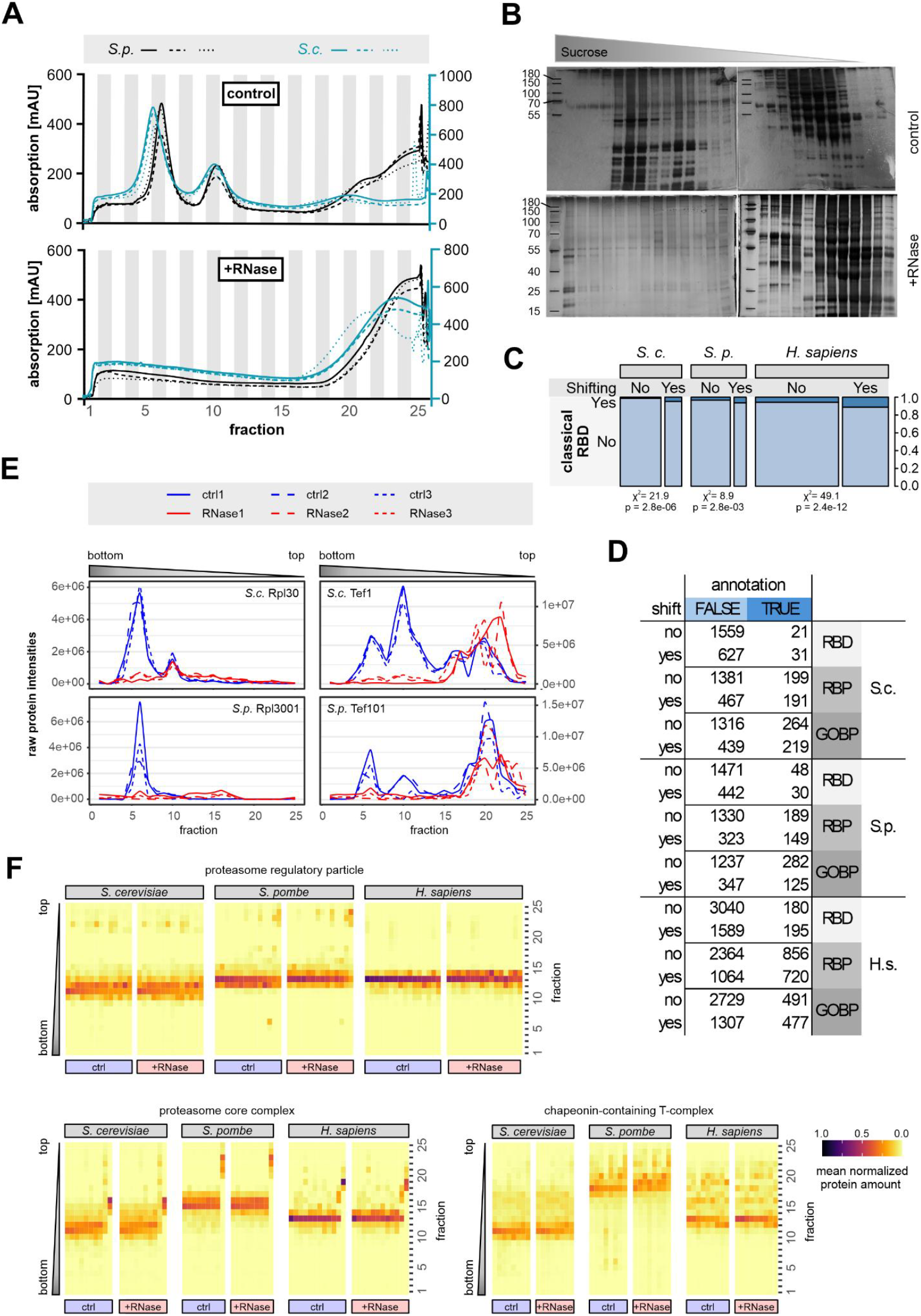
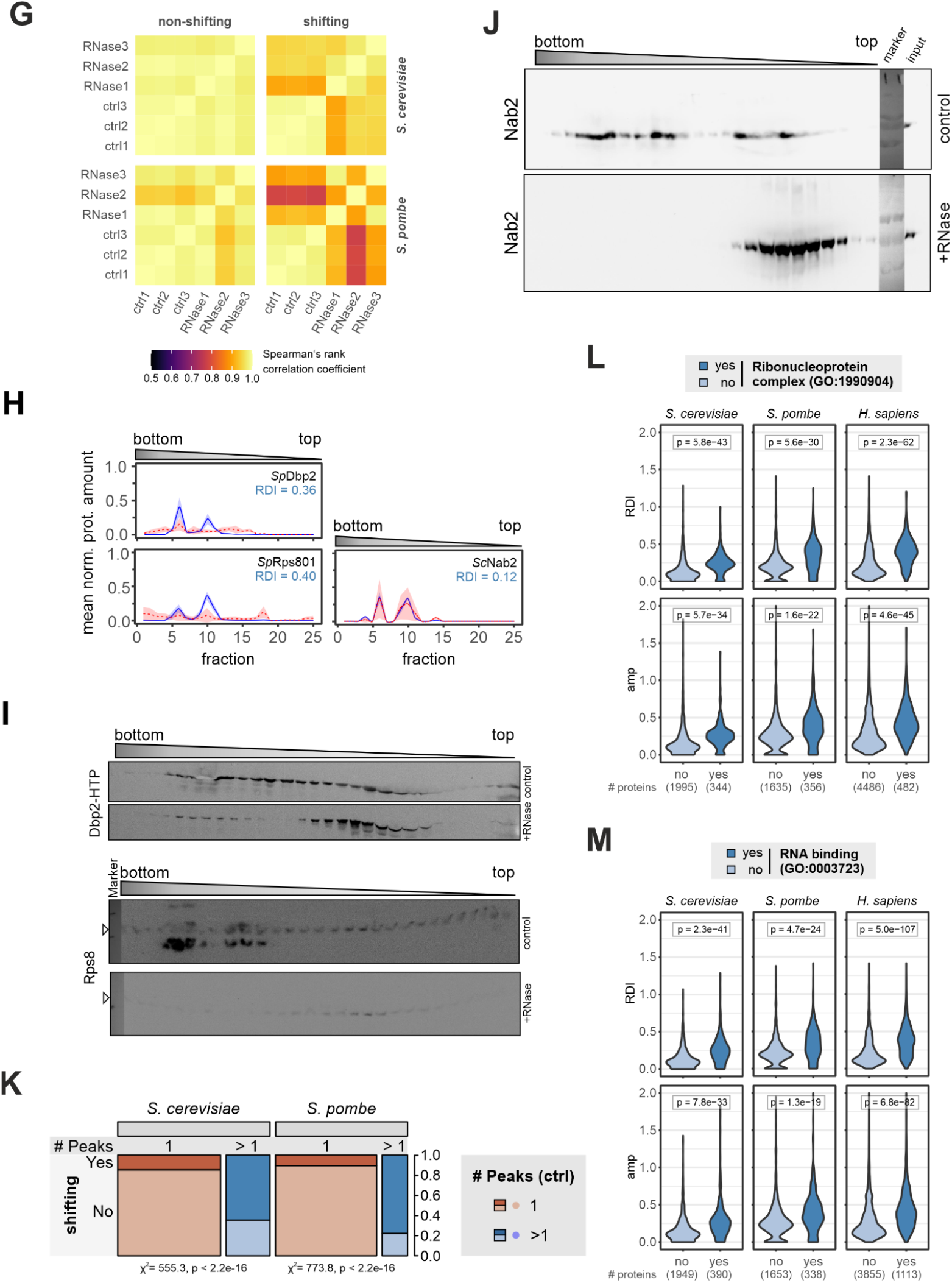
**A** Absorption profile at 280 nm recorded during fractionation of control (top) and RNase-treated gradients (bottom) for *S. pombe* (black) and *S. cerevisiae* replicates (turquoise). The gradients were fractionated on an ÄKTA start at a maximum speed of 1 mL / min. **B** Silver staining of *S. pombe* gradients confirming the loss in protein signal from heavier fractions as detected by A280 nm absorption as in (A). Fractions were TCA-precipitated and resolved in TUS-buffer, then separated on a 12% SDS-Page before silver staining. **C** Proportion of proteins among non-shifting and shifting proteins containing a classical RNA-binding domain (RBD) for *S. cerevisiae* and *S. pombe* compared to the published human data set (Caudron-Herger et al., 2019). Pfam identifiers for classical RBDs were retrieved from Kilchert et al. (2020). Areas are proportional to protein numbers within each category across organisms. Results of Pearson’s Chi-squared test with Yates’ continuity correction are indicated. Absolute protein counts for each category are provided in (D). **D** Protein counts in the indicated categories (RBD: containing classical RNA-binding domain; RBP: RNA binding; GOBP: RNA metabolic process); source data for the spine plots in Figure 1B and (C). **E** Raw protein intensities across the gradients are reproducible between replicates. Raw protein intensities for ribosomal protein L30 (Rpl30/Rpl3001) and translation elongation factor EF-1 alpha (Tef1/Tef101) for the three replicates of RNase-treated (red) and control gradients (blue) from *S. cerevisiae* (left) and *S. pombe* (right) across the 25 fractions from the bottom to the top of the gradient as indicated. **F** Sedimentation behaviour of various multiprotein complexes without known links to RNA metabolism in control and RNase-treated gradients. Components were retrieved from Ensembl using the following GO terms: GO:0005839, GO:0008540, GO:0005832. Human data from Caudron-Herger et al., 2019. **G** Correlation matrices for replicates of the density gradient ultracentrifugation experiment. Heat maps displaying Spearman’s rank correlation between all pairwise comparisons for RNase-treated and control gradients for proteins classified as non-shifting (left) or shifting (right) in *S. cerevisiae* (top) and *S. pombe* lysates (bottom). Correlation coefficients were calculated across all fractions using MS intensities for all proteins detected in at least one of the fractions. Fraction 10 of replicate 3 of the *S. pombe* control gradient contained very little total protein and was disregarded in the analysis. **H** Gradient profiles corresponding to the proteins visualized in the Western blots in (H) and (J): *S. pombe* DEAD-box ATPase Dbp2 (top left) and 40S ribosomal protein S8 (bottom left), and *S. cerevisiae* Nab2 (right) showing mean normalized protein amounts across the gradients for control lysates (blue) and RNase-treated lysates (red). The shaded area behind the curves corresponds to +/-one standard deviation. The RNA dependence index (RDI) was calculated as the Euclidean distance between control and RNase-treated curves, as depicted in 1E. **I** Confirmatory Western blot of control and RNase-treated gradients of *S. pombe* against the well-characterised RNA-binding proteins DEAD-box ATPase Dbp2 (HTP-tagged, PAP-HRP-antibody) and the 40S ribosomal S8 subunit confirms RNA-dependent behaviour. A His-tagged, recombinantly expressed protein from *A. aeolicus* (Aq_880, Nickel et al. 2017) was used as spike-in to clearly assign the signals to the fractions and is marked with an arrowhead. A noticeable shift of Dbp2 to lower sucrose densities is observed following RNase treatment, while 40S ribosomal protein S8, which peaks in heavy fractions in the untreated control, dispersed and was no longer detected after RNase treatment. **J** Western blot of control and RNase-treated gradients of *S. cerevisiae* against the nuclear poly(A)-binding protein Nab2 (mouse IgG1, clone 3F2; M. Swanson lab (Hector et al., 2002)). The Western blot shows a more pronounced shift than the MS data, which showed high variation among RNase-treated gradients. **K** Proportion of proteins peaking at one or multiple positions in the control gradients (brown and blue, respectively) that shift upon RNase treatment. Results of Pearson’s Chi-squared test with Yates’ continuity correction are indicated. **L** Comparison of RNA dependence indices (RDI, top) and shift amplitudes (amp, bottom) for proteins with and without annotation as “ribonucleoprotein complex” component (GO:1990904). Absolute protein counts for each category are provided below the graph. The displayed p-values for the pair-wise comparisons were calculated using a two-sided Wilcoxon rank sum test. Human data from Caudron-Herger et al., 2019. **M** Comparison of RNA dependence indices (RDI, top) and shift amplitudes (amp, bottom) for proteins with and without annotation as “RNA binding” (GO:0003723) as in (M). Human data from Caudron-Herger et al., 2019.

**Supplementary Figure 2:**
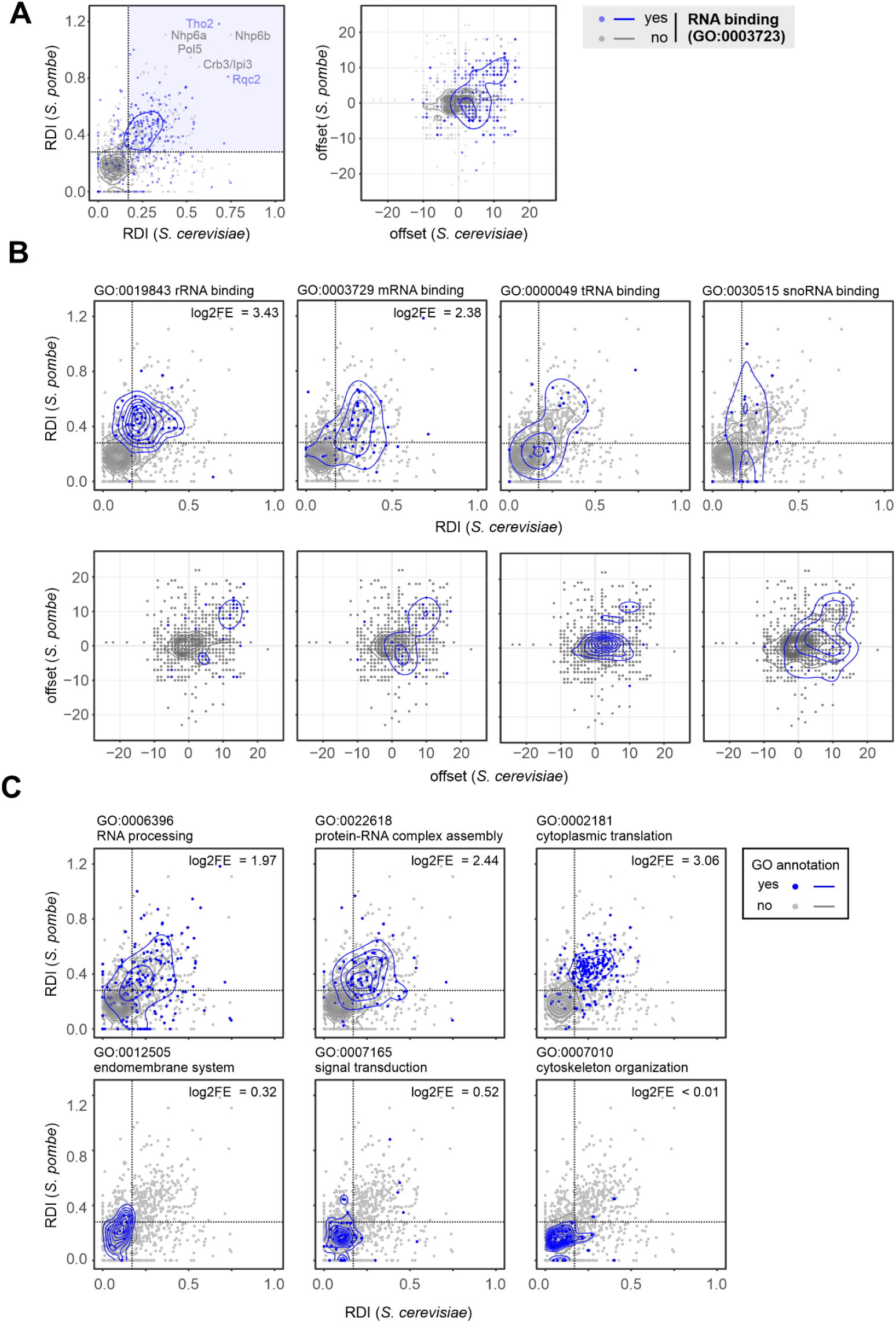
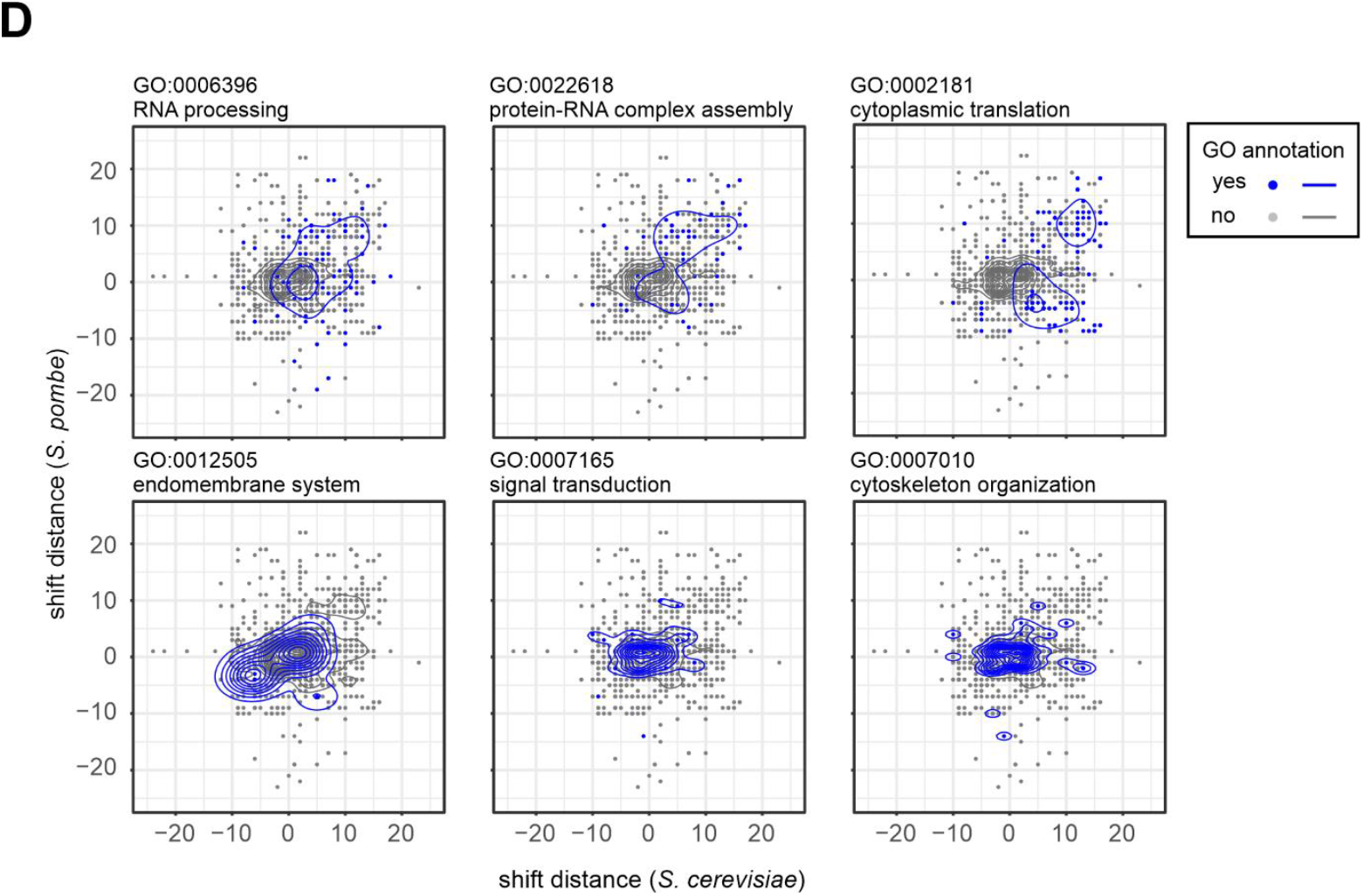
**A** Comparison of the RNA dependence indices (RDI, left panel) and shift offset (right) of orthologous protein pairs in *S. cerevisiae* and *S. pombe*. Proteins annotated as “RNA binding” (GO: 0003723) are shown in blue, all other proteins in grey. Contours of the 2d density estimates for both populations are shown in blue and grey, respectively. Annotations were retrieved for the *S. pombe* orthologues. The identities of the proteins with the highest RDI in both organisms are indicated. The dotted lines represent the cut-offs that were used for GO term enrichment analysis shown in Figure 2B. **B** Comparison of the RNA dependence index (RDI, top panels) and shift distances (bottom panels) of orthologous protein pairs in *S. cerevisiae* and *S. pombe* associated with different classes of RNA (rRNA: ribosomal RNA; mRNA: messenger RNA; tRNA: transfer RNA; snoRNA: small nucleolar RNA). Proteins annotated with the indicated GO term are shown in blue, all other proteins in grey. Contours of the 2d density estimates for both populations are shown in blue and grey, respectively. Annotations were retrieved for the *S. pombe* orthologues. Log2-fold enrichment (log2FE) of annotated proteins in the upper right quadrant relative to their expected number is indicated; proteins annotated as binding to tRNA or snoRNA were not significantly enriched. **C** Comparison of the RDIs of orthologous protein pairs in *S. cerevisiae* and *S. pombe* as in A. Proteins with RNA metabolism-related GO annotations are enriched in the upper right quadrant (top panels) compared to unrelated functional categories (bottom panels). Proteins annotated with the GO terms indicated above the plots are shown in blue. Log2-fold enrichment (log2FE) of annotated proteins in the upper right quadrant relative to their expected number is indicated. **D** Comparison of the shift offset of orthologous proteins in *S. cerevisiae* and *S. pombe* associated with RNA metabolism (top panels) or unrelated pathways (bottom panels) as in (A).

**Supplementary Figure 3:**
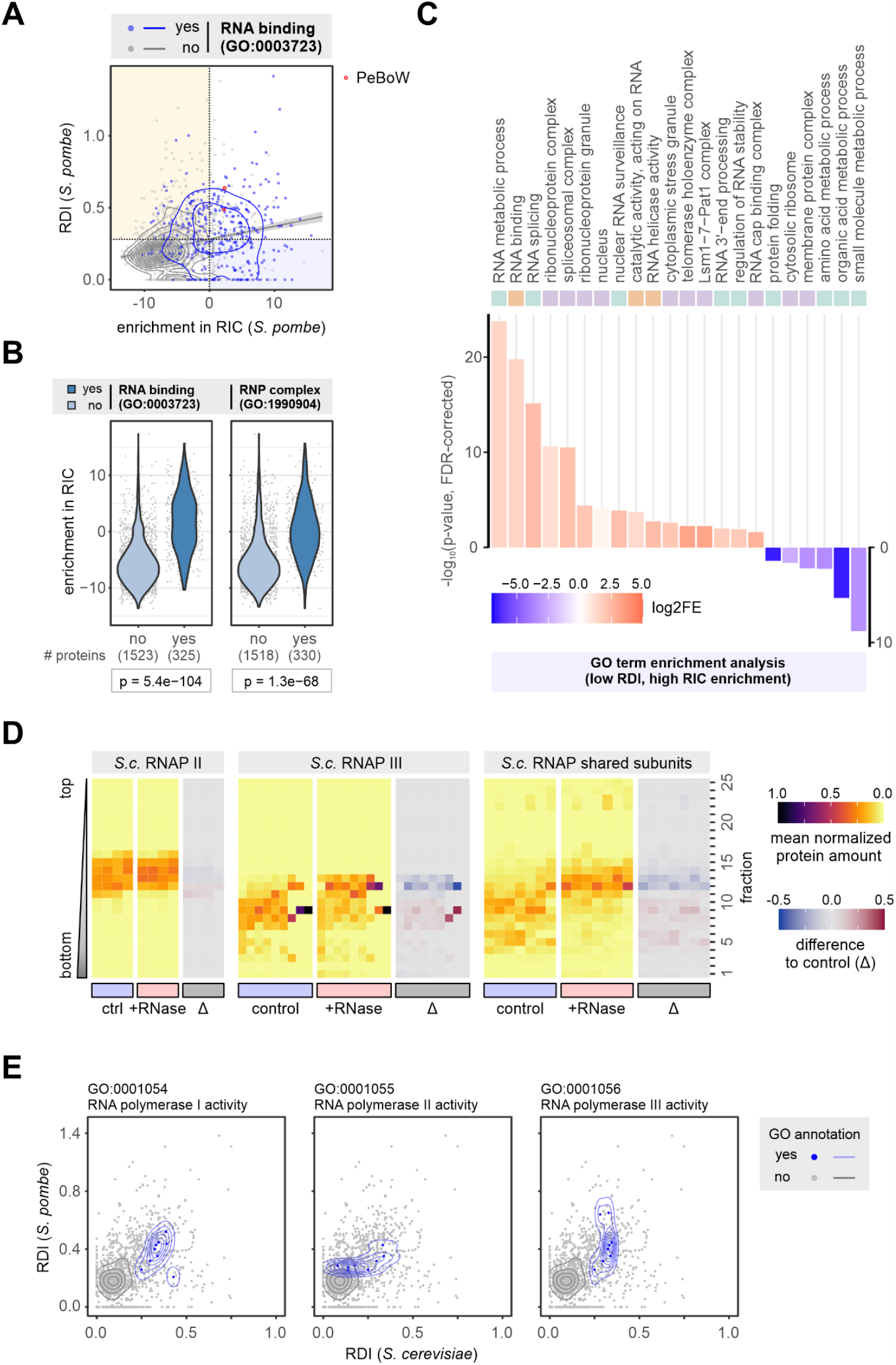
**A** Comparison of RNA independence index (RDI) based on the R-DeeP experiment (this study) and enrichment after poly(A)+ RNA interactome capture (RIC, Kilchert et al. 2020) RIC enrichment is shown as the fold change of mean MS intensities (log2) of proteins recovered from oligo(dT) pull-downs of UV-crosslinked samples relative to the whole-cell extract. Proteins annotated as “RNA binding” (GO:0003723) are shown in blue, all other proteins in grey. Red circles denote PeBoW subunits, one of which was not detected in RIC. **B** Enrichment after poly(A)+ RNA interactome capture for proteins annotated or not as “RNA binding” (GO:0003723; left) or component of “ribonucleoprotein complex” (GO:1990904; right). The displayed p-values for the pair-wise comparisons were calculated using a two-sided Wilcoxon rank sum test. **C** Overrepresented GO terms among proteins with low RDI in *S. pombe* and high enrichment in RIC. GO term enrichment analysis is based on the PANTHER classification system. P-values were calculated using Fisher’s exact test with false discovery rate (FDR) correction. **D** Distribution of RNA polymerase II and III components of *S. cerevisiae* in the gradients. Subunits that are shared between the three RNA polymerases are shown separately. The difference of the mean normalized protein amounts across the RNase-treated gradients to the control is also shown, with signals lost after RNase treatment in red and signals gained in blue. **E** Comparison of the RDI of RNA polymerase I, II and III components in *S. cerevisiae* and *S. pombe*. Proteins annotated with the indicated GO term are shown in blue, all other proteins in grey. Contours of the 2d density estimates for both populations are shown in blue and grey, respectively. Annotations were retrieved for the *S. pombe* orthologues. Shared subunits are included and shown in more than one panel.

**Supplementary Figure 4:**
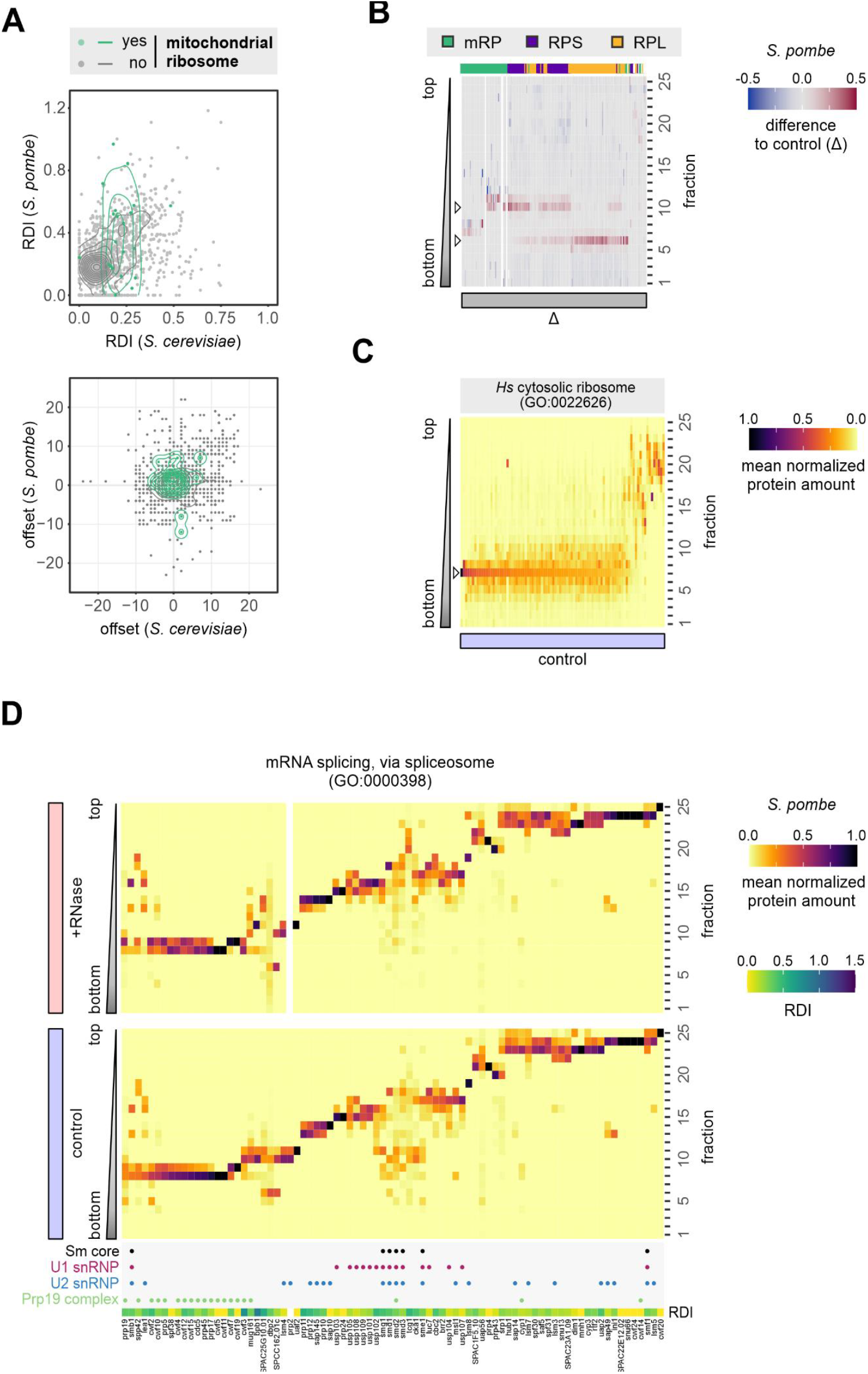
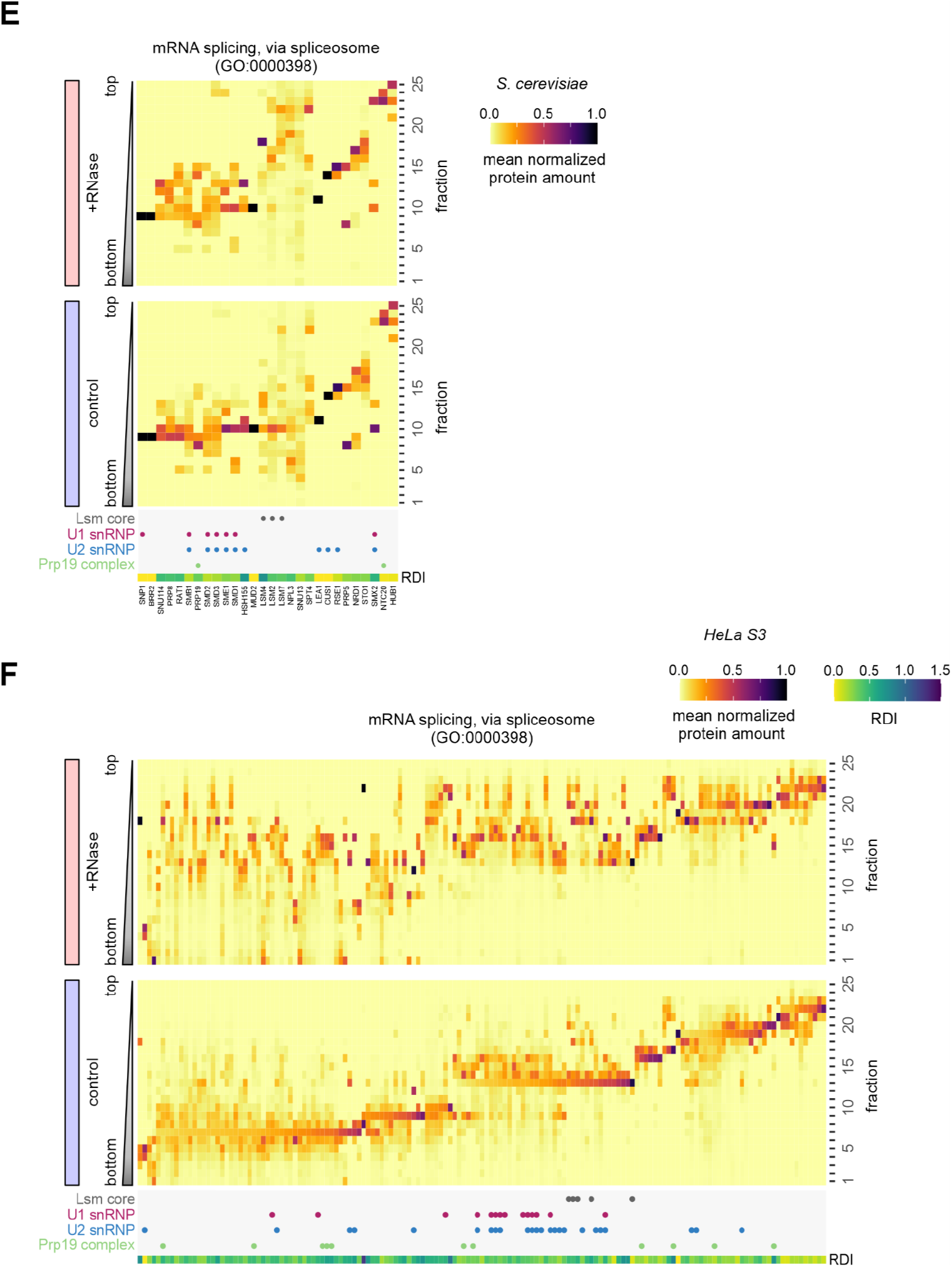
**A** Comparison of the RNA dependence index (RDI, left panel) and shift offset (right panel) of orthologous protein pairs in *S. cerevisiae* and *S. pombe*. Components of the mitochondrial ribosome are shown in green, all other proteins in grey. Contours of the 2d density estimates for both populations are shown in green and grey, respectively. Annotations for GO:0005761 “mitochondrial ribosome” were retrieved from PomBase (Rutherford et al., 2024). **B** Changes to the sedimentation pattern of *S. pombe* ribosomal proteins after RNase treatment shown as difference to the control with signals lost after RNase treatment in red and signals gained in blue. The order reflects 15 clusters that were identified by hierarchical clustering of all ribosomal proteins (GO:0005840) based on mean normalized protein amounts within all fractions of the control gradients as in Figure 3C. Annotations as mitochondrial ribosome (mRP) and the small (RPS) and large (RPL) subunits of the cytoplasmic ribosome are indicated above the plot in green, purple and yellow, respectively. **C** Distribution of ribosomal proteins in HeLa S3 control gradients (Data from Caudron-Herger et al., 2019). **D** Distribution of *S. pombe* proteins involved in mRNA splicing, via spliceosome (GO:0000398) in control gradients (bottom) and RNase-treated gradients (top). Proteins were grouped into 15 clusters identified by hierarchical clustering based on mean normalized protein amounts within all fractions of the control gradients. Both the order of clusters and the order of proteins within clusters were adjusted for clarity of visualization. For each protein, the RDI and association with selected complexes is indicated below the plot. **E** As in (D), but for *S. cerevisiae*. **F** As in (D), but for human proteins (HeLa S3 gradient data from Caudron-Herger et al., 2019).

**Supplementary Figure S5:**
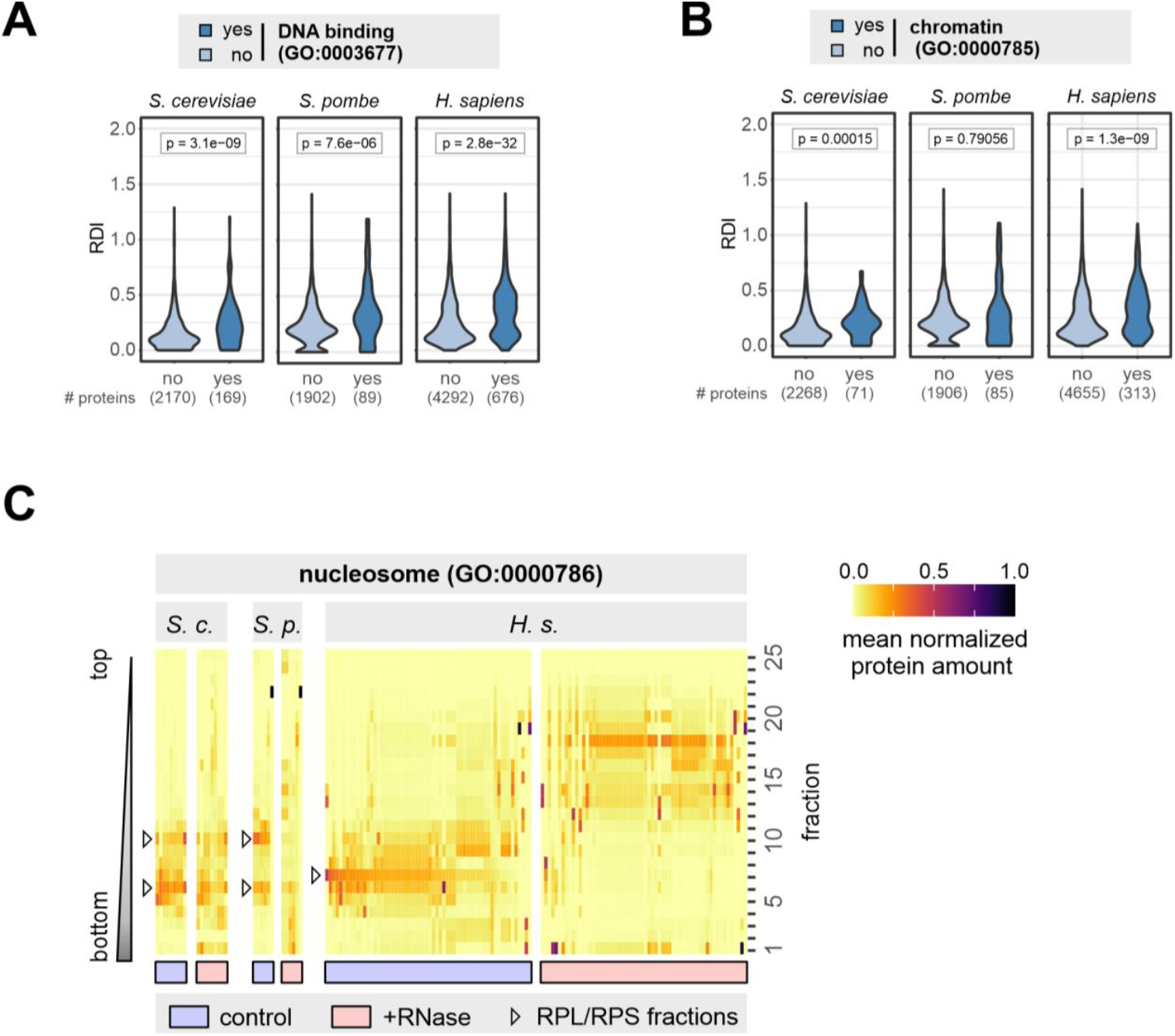
**A** RDIs for proteins with and without annotation as “DNA binding” (GO:0003766). The displayed p-values for the pair-wise comparisons were calculated using a two-sided Wilcoxon rank sum test. Human data from Caudron-Herger et al., 2019. **B** RDIs for proteins with and without annotation as component of “chromatin” (GO:0000785). The displayed p-values for the pair-wise comparisons were calculated using a two-sided Wilcoxon rank sum test. Human data from Caudron-Herger et al., 2019. **C** Distribution of nucleosome components (GO: 0000786) in control and RNase-treated gradients. The fractions in which ribosomal proteins of the small or large subunit preferentially sediment are indicated by arrowhead (compare Figure 4C and S4C). Human data from Caudron-Herger et al. 2019.

